# Complete Chloroplast Genomes of *Saccharum giganteum*, *Saccharum longisetosum*, *Cleistachne sorghoides, Sarga timorense, Narenga porphyrocoma* and *Tripsacum dactyloides*. Comparisons with ITS phylogeny and Placement within *Saccharum*

**DOI:** 10.1101/2020.06.12.149476

**Authors:** Dyfed Lloyd Evans, Ben Hughes

## Abstract

The first complete chloroplast and Internal Transcribed Sequence (ITS) cassette sequences for the species: *Saccharum giganteum*, *Saccharum longisetosum*, *Cleistachne sorghoides, Saccharum narenga* and *Tripsacum dactyloides* are presented. Corresponding sequences for a new isolate of *Sarga timorense* were assembled. Phylogenetic analyses place *S. giganteum*, *S. longisetosum* and *S. narenga* within the Saccharinae but distinct from Saccharum, whilst *C. sorghoides* emerges as a member of genus *Sarga* and *Tripsacum datyloides* as a member of the Tripsacinae. Comparison of chloroplast and ITS phylogenies reveal complex reticulate evolution within the Saccharinae, with *S. giganteum*, *S. longisetosum* and *S. narenga*, despite having the same base chromosome count (15) having different evolutionary origins; making them members of different genera and not members of genus *Saccharum*. The importance of reticulate evolution in the origins of Andropogoneae, particularly the Saccharinae and the unique positions of *Saccharum* and *Miscanthus* is discussed.

## Introduction

Despite recent advances in plant low copy number gene analysis (Zeng et al. 2014), combined whole chloroplast and low copy number gene analyses (Lloyd Evans et al. 2019) the phylogenetic placement of many species related to sugarcane remain uncertain. This is partly due to lack of taxon sampling and partly due to many of these species having been lumped into genus *Saccharum* with no molecular phylogenetic verification. Moreover, the only large-scale analyses to include many of these species were performed over a decade ago (Hodkinson et al. 2002). Indeed, there is no complete chloroplast sequence for any member of the New World Erianthus species (particularly the type species *Erianthus giganteus* (syn *Saccharum giganteum*) or the trans-Himalayan species of saccharum/miscanthus (as exemplified by *Erianthus rockii* (syn *Saccharum longisetosum*)) currently deposited in ENA/GenBank. Our recent study (Lloyd Evans et al. 2019) removed the Old-World species from genus *Erianthus*, placing them within their own genus, *Tripidium*. This has made the examination of the taxonomic placement of *Erianthus giganteus* (*Saccharum giganteum*) all the more urgent. Particularly as members of genus *Tripidum* (under the guise of *Erianthus* sect *Ripidium*) have been used in introgression breeding with Sugarcane (typically without much success) (Piperidis et al. 2000). However, the potentially more closely related New World *Erianthus* species have not been employed within sugarcane breeding programmes.

*Erianthus gianteus* is the type species for genus *Erianthus* as a whole. The species was first described as: *Anthoxanthum giganteum* by Thomas Walter in 1788 (Walter, 1788). It was re-classified as *Erianthus giganteus* by A.M.F.J Palisot de Beauvois in an essay of 1812 (Palisot de Beauvois, 1812). This is important, as apart from *Saccharum officinarum* and *Sccharum spontaneum* (both defined by Linnaeus in his first volume of *Species Plantarum*, 1753) (Linnaeus, 1753) the definition of *Erianthus* antedates the other genera most closely related to Saccharum. Accurate phylogenomic placement of *Erianthus sensu stricto* (in the strict sense) could have a marked impact on the naming of genera closely related to *Saccharum*. However, to complicate matters taxonomically, Christian Hendrik Persoon defined the species as *Saccharum giganteum* in 1805 (Persoon 1805).

In terms of nomenclature, Kew’s GrassBase (Clayton et al. 2002) presents *Saccharum giganteum* as the accepted name, whilst Tropicos (http://tropicos.org/) presents both *Erianthus giganteus* and *Saccharum giganteum* as accepted names.

There is a similar confusion with the second species chosen for this study, *Erianthus rockii* (the name typically employed by sugarcane breeders), as described by YL Keng in 1939 (Keng, 1939). However, NJ Andersson first described the species as *Erianthus longisetosus* in 1855 (Andersson 1855). V. Narayanaswami later incorporated the species into genus Saccahrum as *Saccharum longisetosum* in 1940 (Narayanaswami 1940).

George Bentham first described *Cleistachne sorghoides* in 1882 (Bentham 1882). Tropicos gives *Cleistachne sorghoides* as the accepted name, as does Kew’s Grassbase, but the NCBI Taxonomy (https://www.ncbi.nlm.nih.gov/taxonomy) gives *Sorghum sorghoides* (Benth.) Q.Liu & P.M.Peterson as the accepted name, placing this species within genus *Sorghum*. Our previous analysis, using extended ITS sequences (Snyman et al, 2108), placed *Cleistachne sorghoide*s within genus *Sarga* making it distinct from Sorghum. However, no complete chloroplast genome sequence for this species was available for direct comparison.

*Narenga porphyrocoma* (Bor) hybridizes readily with sugarcane (Mukherjee 1957; Liu et al. 2012), as such it could either be a close relative of *Saccharum* or it could be a member of *Saccharum* (indeed its currently accepted name is *Saccharum narenga* (Nees) Wall. ex Hack.). For this study, both the Chloroplast genome and the extended ITS sequence of *Narenga porphyracoma* were sequenced.

The Tripsacinae, a subtribe that includes *Zea mays* L., is a very useful outgroup for phylogenetic analyses of the core Andropogoneae, as it lies distal to the core Andropogoneae, the subtribe that delimits those members of the Andropogoneae that can be part of the Saccharinae (sugarcane’s close relatives) and those species or genera that cannot be part of the Saccharinae (Kellogg 2013). It is therefore surprising that the complete chloroplast of one of maize’s more distal relatives, *Tripsacum dactyloides* (gammagrass) has not previously been sequenced and assembled. For phylogenomic analyses the complete chloroplast has been mooted as a ‘super barcode’ for plants, particularly grasses (Krawczyk et al. 2018). The relative ease by which chloroplast DNA can be isolated and assembled from whole genome short read data or sequenced from PCR amplification from whole plant DNA isolates has made chloroplast genome assembly and phylogenomics much simpler. However, it has recently become clear (Connor 2004; Kellogg et al. 1996; Hinsinger et al. 2014; Záveská et al. 2016; Folk et al. 2017) that plant evolution is much more complex than previously realised and that reticulate (network) evolution is much more common than previously thought. This is particularly the case in the Andropogoneae subtribe of grasses, where ancestral cross species hybridization has been very common (Estep et al. 2014, Lloyd Evans et al. 2019). As a result, the interpretation of the ‘true’ evolutionary origin of any species needs a comparison of both the plastid and nuclear genome evolution. The effect of such ancestral hybridizations on speciation also warrants further investigation.

The internally transcribed sequences of the ribosomal RNA cassette have traditionally been used to determine plant phylogenetic relationships (Hodkinson et al. 2002). ITS regions are very common within the plant genome, being expressed at high transcript levels, and therefore are very easy to amplify and sequence. However, traditional ITS regions of 600bp to 800bp provide too little phylogenetic signal for reliable phylogenies (branches of closely-related species typically collapse to polytomies) (Álvarez and Wendel 2003). Recently we demonstrated the utility of longer ITS regions for generating more robust nuclear-based phylogenies in the Andropogoneae (Snyman et al. 2018). In this study, near complete ITS cassettes (~7500bp) were sequenced from the target species of this study along with sugarcane cultivar SP80-3280. These regions have recently come to attention as a second super barcode for plant phylogenetics (Chen et al. 2018).

Cytogenetic studies also present invaluable information about chromosome numbers and potential chromosomal incompatibilities between *Saccharum* species and genera closely related to *Saccharum*. *Saccharum* itself has a base chromosome number (x) of 10 (Li et al. 1959), though chromosomal fusion has reduced this to x=8 in *Saccharum spontaneum* (Ha et al. 1999). *Sorghum* also has a base chromosome number of 10, though *Sarga* has a base chromosome number of five (Gu et al. 1984). A genome doubling and chromosomal fusion in *Miscanthus* led to a base chromosome number of 19 (Adati 1958). African *Miscanthidium* species have a base chromosome number of 15 (Strydom et al. 2000), as do New World *Erianthus* species (eg *Erianthus giganteus*) (Jensen et al. 1989). *Erianthus longisetosus* (syn *Erianthus rockii*) has two reports with a diploid count of 60, yielding a monoploid count of 30 and a base count of 15; thus *E. rockii* is a tetraploid (Burner 1991). A further report from India (Löve 1976) gives a haploid and base count of n = 15 (x=15) for this species, indicating that the sample analyzed is diploid. We therefore have *E. rockii* (*E. longisetosus*) with different ploidy levels in the wild, but all having a base chromosome number of 15. *Narenga porphyrocoma* also has a base chromosome count of 15 (Sreenivasan and Sreenivasan 1984). Thus, chromosome counts would indicate that there is a clear division between *Sorghum*, *Sarga*, *Miscanthus* and *Saccharum*. *Erianthus*, *Miscanthidium* and *Narenga* have the same chromosome counts, but no analysis to date has examined the degree of their inter-relatedness.

Interestingly, *Chleistachne sorghoides* lies outside this series with a base chromosome number of nine (Celarier 1958), which could have arisen as a chromosome fusion in the 10 base chromosomes of sorghum or by the result of a genome duplication and chromosomal fusion in *Sarga*. The phylogenomic placement of *Cleistachne sorghoides* would answer this puzzle.

To maximize sequencing efficiency (and reduce costs), chloroplasts and complete ITS regions were isolated from total DNA and RNA, respectively, by magnetic bead probe baiting before being sequenced as long reads with Oxford Nanopore Technologies (ONT) MinION sequencing technology.

## Materials and Methods

### Plant materials

*Chleistachne sorghoides* was collected near the border of South Africa, Swaziland and Mozambique (just outside Mbuzini; geolocation: −25.934259,31.942466) and *Sarga timorense* seeds were collected outside Soe, South Timor (geolocation: – 9.880422,124.283087). Seeds for *Tripsacum dactyloides* inbred line TDD39103 were kindly donated by the Cold Spring Harbor Laboratory. Seeds for *Erianthus giganteus* cv Mountain Sunset were purchased from Hoffman Nursery, Rougemont, NC. Seeds for *Erianthus rockii* (from a specimen originally collected outside Nuijang, Yunan Province, China) were obtained from B&T World Seeds, Aigue Vives, France. Leaf material for *Narenga porphyrocoma* was sourced non-destructively in the wild (found on the outskirts of Teluk Intan, Perak, Malaysia (geolocation: 4.003058, 101.048456)).

Seeds were laboratory grown under optimal conditions (heat, day length, humidity) to 6cm tall prior to harvesting. In all cases, six plantlets were allowed to grow to maturity (flowering) to confirm the identity of each species.

### DNA Isolation

Wherever possible young tissue was snap frozen in liquid nitrogen prior to DNA extraction. Whole plantlet or leaf tissue (5g) was ground in a pestle and mortar under liquid nitrogen and ground tissue was immediately utilized for DNA extraction using the modified CTAB method described by Doyle and Doyle (1987). DNA quality was determined spectrophotometrically by assessing the purity using the measurement of absorbance ratio, A260/A280, with a plate reader.

### RNA Isolation

Total RNA was isolated from whole plantlet or leaf tissue (5g) that was ground in a pestle and mortar under liquid nitrogen. Ground tissue was immediately utilized for RNA isolation with the PureLink RNA Mini Kit (Thermo Fisher, Altrincham, UK) according to the manufacturer’s protocols.

### Magnetic Bead Capture Primer DNA Isolation

For chloroplast capture, 30bp capture primers (T_m_ between 63-84°C) were developed based on tRNA genes (these are highly conserved across evolutionary time). The chloroplasts of *Saccharum* hybrid cv SP80-3280, *Miscathus sinenis* cv Andante, Sorghum bicolor cv BTx632 and Zea mays cv B73 were aligned, tRNAs were mapped and conserved 30-mer sequences from 15 tRNA genes covering the entire chloroplast (Table 1) were designed. For ITS capture five 35-mer primers (to account for differences between DNA and RNA) were designed covering 2x 18s, 1x 5.8s and 2x 25s sites (Table 1). Capture primers were designed with a 10bp inosine 3’ extension attached to streptavidin. Primers were tested and validated using in-silico PCR with Amplify4 (Engels 2015). Two additional blocking primers were developed to prevent cp genome self-annealing from displacing the capture primers (see Shepard and Rae 1997). The biotin-bound capture primers (Sigma-Aldrich, Gillingham, UK) were attached to biotinylated magnetic beads (Promega, Southampton, UK) were washed in 100μl 6x SSC, captured with a magnetic strand (Promega) and washed again. This process was repeated three times. The beads were resuspended in 100μl bead block buffer (2% I-Block, Thermo Fisher) and 0.5% SDS in PBS buffer. The solution was incubated at room temperature for 10 minutes. Pre-treated beads were captured with a magnetic strand and re-suspended in 100μl 6x SSC prior to mixing with the biotinylated probes (one set for the CP probes and one set for the ITS probes). Beads and probes were incubated at room temperature for 10 mins with agitation by gentle flicking. Bound bead and probe complexes were washed at room temperature with four washes of 6x SSC and bead capture with the magnetic strand. The probe-bound beads were combined with isolated DNA/RNA (separate aliquots for chloroplast and ITS RNA), and for CP genome capture the additional blocking primers in 6x SSC buffer and were incubated at 95°C for 10 min prior to annealing for 60 min at 60°C (for RNA ITS capture 5μl RNA preservation buffer [25mM sodium citrate, 10 mM EDTA, 0.2M guanidium thiocyanate, 79g ammonium sulfate/100ml solution, adjust to pH 5.2 with 1M sulfuric acid] was added to the SSC in each use. A magnetic strand (Promega) was employed to draw aside the beads. The beads were subject to three washes of 200μl 1x SSC prior to final capture with the magnetic strip. To release the bound DNA the beads were re-suspended in 50μl H2O and heated to 75°C for 5 min. Whilst still hot, the magnetic strand was used to remove the magnetic beads. As this methodology captures both strands of CP genome DNA, as the DNA solution was allowed to cool double stranded DNA re-annealed. Beads were re-used and were washed at 1mM NaCl and 95°C for 10 mins between each use.

**Table 1.**
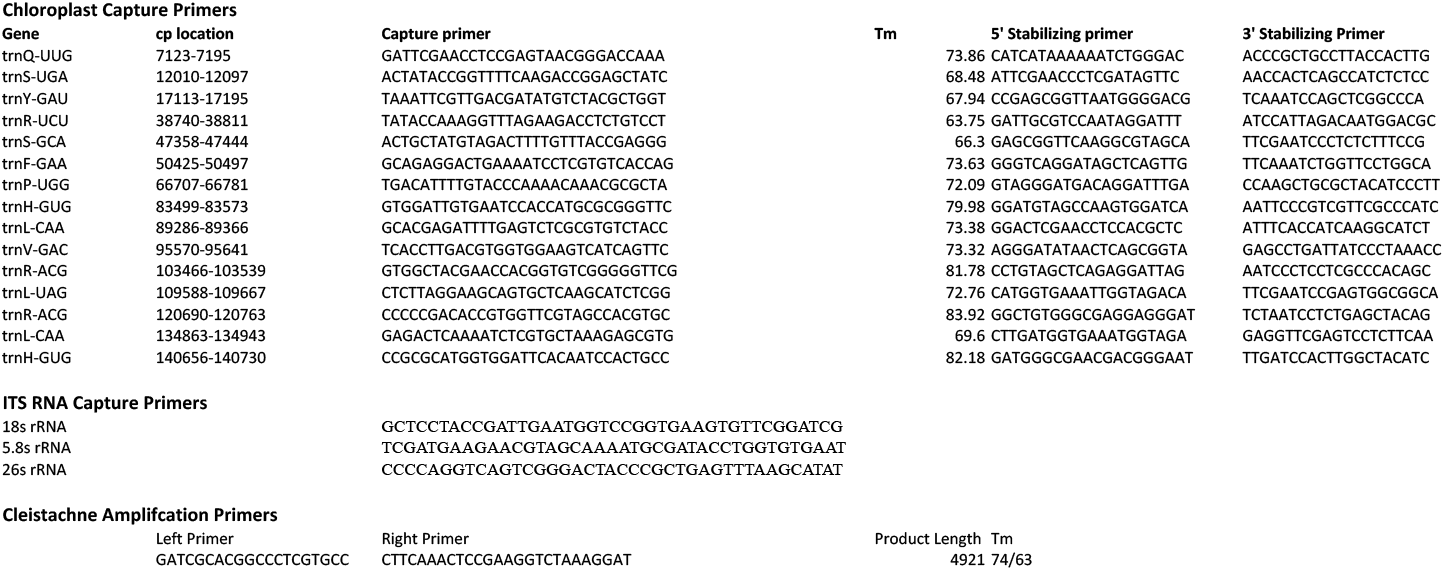
List of capture primers developed for whole chloroplast and complete ITS DNA isolation. A complete list of the capture primers developed for isolating whole chloroplast Genomes and complet ITS cassette regions in *Cleistachne sorghoides*, Sarga timorense, *Erianthus longisetosus*, *Erianthus giganteus*, *Tripsacum dactyloides* and Narenga porphyrcoma. ITS primers were employed for Miscanthus ITS isolation. Due to some DNA degradation, PCR amplification was employed for Cleistachne ITS and these were based on previously assembled *Sarga* sequences (Snyman et al. 2018).

### PCR Primer Design for *Cleistachne sorghoides* ITS

Due to the presence of RNA degradation, PCR amplification of genomic DNA was employed for *Cleistachne* ITS (Table 1). Primers were developed with NCBI’s Primer Design tool (Ye et al. 2012) and were based on previously assembled *Sarga* sequences (Snyman et al. 2018). Primers were tested using in-silico PCR with Amplify4. Amplicons were purified by HPLC column chromatography, concentrated by lyophilization and used directly for MinION sequencing.

### Sequencing and Assembly

Isolated DNA was HPLC size sorted on Phenomenex Luna 5μm 100Å column with endcapping (Phenomenex, Macclesfield, UK) to eliminate any genomic DNA contamination. Eluted DNA was concentrated by lyophilization and prepared for For MinION DNA sequencing, sample preparation was carried out using the Genomic DNA Sequencing Kit SQK-MAP-006 (Oxford Nanopore Technologies) following the manufacturer’s instructions, including the optional NEBNext FFPE DNA repair step (NEB). For RNA (ITS) sequences the Direct RNA sequencing kit (SQK-RNA002) was employed. For sequencing, a 6μl aliquot of pre-sequencing mix was combined with 4 μl Fuel Mix (Oxford Nanopore), 75μl running buffer (Oxford Nanopore) and 66μl water and added to the flow cell. The 48h genomic DNA sequencing script was run in MinKNOW V0.50.2.15 using the 006 workflow. Metrichor V2.33.1 was used for base calling. Raw data from a single MinION flow cell was initially analysed using Metrichor and binned into ‘pass’ or ‘fail’ based on a threshold set at approximately 85 % accuracy (Q9). In our hands, a 12-hour MinION run yielded 90% of the highest quality reads, thus runs were terminated after 12 hours. Reads were separated according to barcode and barcode sequences were removed. Each amplicon was assembled individually with Canu (Koren et al. 2017), using Canu’s in-built error correction.

### Assembly Finishing and Annotation

Canu assemblies emerge as error corrected and polished. In all cases extended ITS assemblies yielded a single sequence. Chloroplast assemblies yielded two sequences, with at least 2kbp overlaps indicating circularity. The two chloroplast isoforms differed only in the orientations of the SSC regions.

As the Andropogoneae are well represented in the Verdant database, chloroplast genomes were initially annotated using the Verdant webserver (http://verdant.iplantcollaborative.org/) by submitting the fully assembled chloroplast as a fasta format file (McKain et al. 2017). Annotation was downloaded in Verdant native (.tbl) format. Additional genes not present in the Verdant system, as identified by previous analyses (Lloyd Evans and Joshi 2016, Lloyd Evans et al. 2019) were mapped using BLAST. Using a custom Perl script (available from https://github.com/gwydion1/bifo-scripts) Verdant annotation, additional annotation, FASTA sequence and NCBI taxonomy files were merged to yield an EMBL-format flatfile which was submitted to ENA. Verdant cannot annotate the inverted SSC regions in the chloroplast isoforms and these were annotated manually based on the annotation of the forward SSC regions. The final annotated assemblies were submitted to the European Nucleotide Archive (ENA) under the project accession: PRJEB31602.

ITS assemblies were annotated against the reference sequence of *Lecomtella madagascariensis* (GenBank: HG315108) (Besnard et al. 2013) ITS and a custom PERL script was employed to convert Blast features into an ENA compatible file format. Annotated ITS assemblies were submitted to the ENA under the project accession: PRJEB31603.

### Assembly of Additional ITS regions

As full length ITS sequences are not yet common in the global sequence databases, the NCBI sequence read archive (SRA) was mined for potentially assemblable datasets in the Andropogoneae. The following species and associated SRA datasets were identified: *Tripidium procerum* (SRR2891248); *Tripidium rufipilum* (SRR2891271); *Themeda triandra* (SRR7529014); *Cymbopogon nardus* (ERR2040774); *Cymbopogon ciratus* (SRR11229723); *Cymbopogon wintriantus* (SRR1614278), *Hyparrhenia rufa* (SRR7121600); *Sorghum amplum* (SRR2094869-SRR2094876); *Sorghum propinquum* (SRR999028); *Sorghum halepense* (SRR486216); *Sorghum bicolor* (SRR5271056); *Sorghum arundinaceum* (SRR999026); *Miscanthus sacchariflorus* cv Hercules (SRR486749); *Miscanthus floridulus* (SRR10988838); *Saccharum spontaneum* SES196 (SRR2899231); *Saccharum robustum* NG57-054 (SRR2899233); *Miscanthidium junceum* (SRR396848); *Sarga brachypodium* (SRR3938607); *Sarga versicolor* (SRR427175); *Sarga plumosum* (SRR8666235); *Coix lacryma-jobi* (SRR10208265); Coix *aquatica* (SRR1634981); *Microstegium vimineum* (ERR2040772) and *Zea luxurians* (SRR5466389).

For assembly, reads were baited with mirabait against a reference, as described previously. If no full-length template available, the longest ITS template from GenBank was used. Reads were assembled with SPAdes using a k-mer series up to 127 if reads were longer than 150bp or a kmer that was the closest odd number below the read length if read lengths were 125bp or less. If the longest assembled contig was less than 7500bp the longest contig was used for baiting and re-assembly. This re-assembly process was continued until the longest contig was 7500bp or longer. Finished assemblies were submitted to Zenodo (as these are third party data). In addition, the full length ITSes of *Saccharum perrieri* (GenBank: MN342165.1) and *Lasiorhachis hildebrandtii* (GenBank: MN342164.1) were added to the dataset.

### Sequence Alignments

Assembled chloroplast genomes were oriented to the same direction and start position before being combined with 43 additional chloroplast genomes from NCBI (see Supplementary Table 1 for a full list) and aligned with SATÉ 2.2.2 (Liu et al., 2009) using default options and the RAxML GTRGAMMA model. Alignments were optimized with Prank (an indel-aware re-aligner) (Löytynoja et al. 2012). Terminal taxa representing well-supported groups as defined by the SATÉ RAxML phylogram were constrained using Prank’s ‘group’ functions. For the chloroplast alignment, a single round of SATÉ followed by Prank yielded an optimal alignment (which was corrected by hand to reduce inserted stretches of more than 20 nucleotides due to a single species only to 10nt).

An equivalent process was employed to align the ITS regions. However, as the ribosomal RNA sequences tend to be well conserved and the ITS regions are more plastid, six rounds of Prank re-alignment and RAxML (version 8.1.17) (Stamakis, 2006) tree topology analyses were required to yield an optimal alignment and a stable phylogeny.

### Phylogenomic Analyses

As determined previously (Lloyd Evans et al. 2019), the following partitioning schema yields optimal results for the Andropogoneae: the whole plastid alignment was divided into LSC, IR and SSC partitions. These partitions were further divided into protein-coding gene, RNA-coding gene and non-coding regions. The regions were isolated with the BeforePhylo.pl (Zhou 2014) script and merged into separate partitions. The IRA region contained only a single tRNA encoding gene, which was added to the SSC RNA-gene partition. This yielded a total of eight partitions. Best-fit evolutionary models for each partition were selected using JModelTest2 (Darriba et al. 2012) and the AICc criterion. The best-fit models were as follows: LSC protein coding: TPM1uf+I+Γ, LSC RNA genes: TVM+Γ; LSC non-coding: TVM+Γ; IRA protein coding: TVM+I+Γ; IRA non-coding: TVM+Γ; SSC protein coding: TPM1uf+I+Γ; SSC RNA-gene: TrN+I+Γ and SSC non-coding: TVM+I+Γ. The partitions determined above and their closest model equivalents were used for all subsequent analyses.

The ITS alignment was divided into rrn18, ITS1, ITS2, rrn5.8 and rrn28 partitons. JModelTest analyses revealed the best fit models to be: 18s RNA: TVM + Γ; ITS1: TPM3uf + Γ; ITS2: TPM3uf + Γ; 5.8s rRNA: JC + Γ; ITS2:GTR+ Γ; 28srRNA:GTR+ Γ. These partitions were employed for all subsequent analyses. Bayesian Inference analyses were run with MrBayes (version 3.1.2) (Ronquist and Huelsenbeck, 2003), Maximum Likelihood analyses were run with IQ-Tree (Nguyen et al. 2015) and single branch test SH-aLRT analyses were run with IQ-Tree (Nguyen et al. 2015).

For both the whole chloroplast and ITS datasets Bayesian Markov Chain Monte Carlo (MCMC) analyses were run with MrBayes 3.1.2, using four chains (3 heated and 1 cold) with default priors run for 20 000 000 generations with sampling every 100th tree. Two separate MrBayes analyses, each of two independent runs, were conducted. To avoid any potential over-partitioning of the data, the posterior distributions and associated parameter variables were monitored for each partition using Tracer v 1.6 (Rambaut et al. 2017). High variance and low effective sample sizes were used as signatures of over-sampling. Burn-in was determined by topological convergence and was judged to be sufficient when the average standard deviation of split frequencies was <0.001 along with the use of the Cumulative and Compare functions of AWTY (Nylander et al. 2008). The first 5 000 000 (25%) sampled generations were discarded as burn-in, and the resultant tree samples were mapped onto the reference phylogram (as determined by maximum likelihood analysis) with the SumTrees 4.0.0 script of the Dendropy 4.0.2 package (Sukumaran and Holder 2010).

Non-parametric bootstrap and single branch SH-aLRT analyses for 5 000 replicates were run with IQ-Tree (Nguyen et al. 2015). Alignments and reference Maximum Likelihood phylogenies are available from Zenodo (Lloyd Evans 2020).

## Results

As expected, being separated by only ~7.5 million years of evolution (Lloyd Evans and Joshi 2106; Lloyd Evans et al 2019), the chloroplast genomes assembled in this study are very similar, ranging in size from 141009 bp in *Saccharum longisetosum* to 1411073 bp in *Saccharum giganteum* (Figure 1). They contained 84 protein coding genes, 30 tRNA genes and 3 ribosomal RNA genes (excluding duplicate copies on the inverted repeats). Each chromosome had a quadripartite structure, with large single copy (LSC), short single copy (SSC) and two inverted repeat (IR_B_, IR_A_) regions. Images of the four chloroplasts derived from genomic sequence are shown in Figure 1. In all cases, as described by Wang and Lanfear, 2019, two isoforms of the chloroplast genome were obtained, differing only by the inversion of the SSC region. Direct comparisons of the chloroplast types can be seen in Supplementary Document 1. Ratios of canonical and inverted SSCs as determined by the Cp-hap pipeline (Wang and Lanfear, 2019) were as follows: *Chleistachne sorghoides* 51.4:48.6; *Sarga timorense* 51.1:48.9; *Tripsacum dactyloides* 50.4:49.6; *Erianthus giganteus* 53.3:46.7; *Erianthus rockii* 53.1:46.9 and *Narenga porphyrocoma* 47.2:52.8.

**Figure 1.**
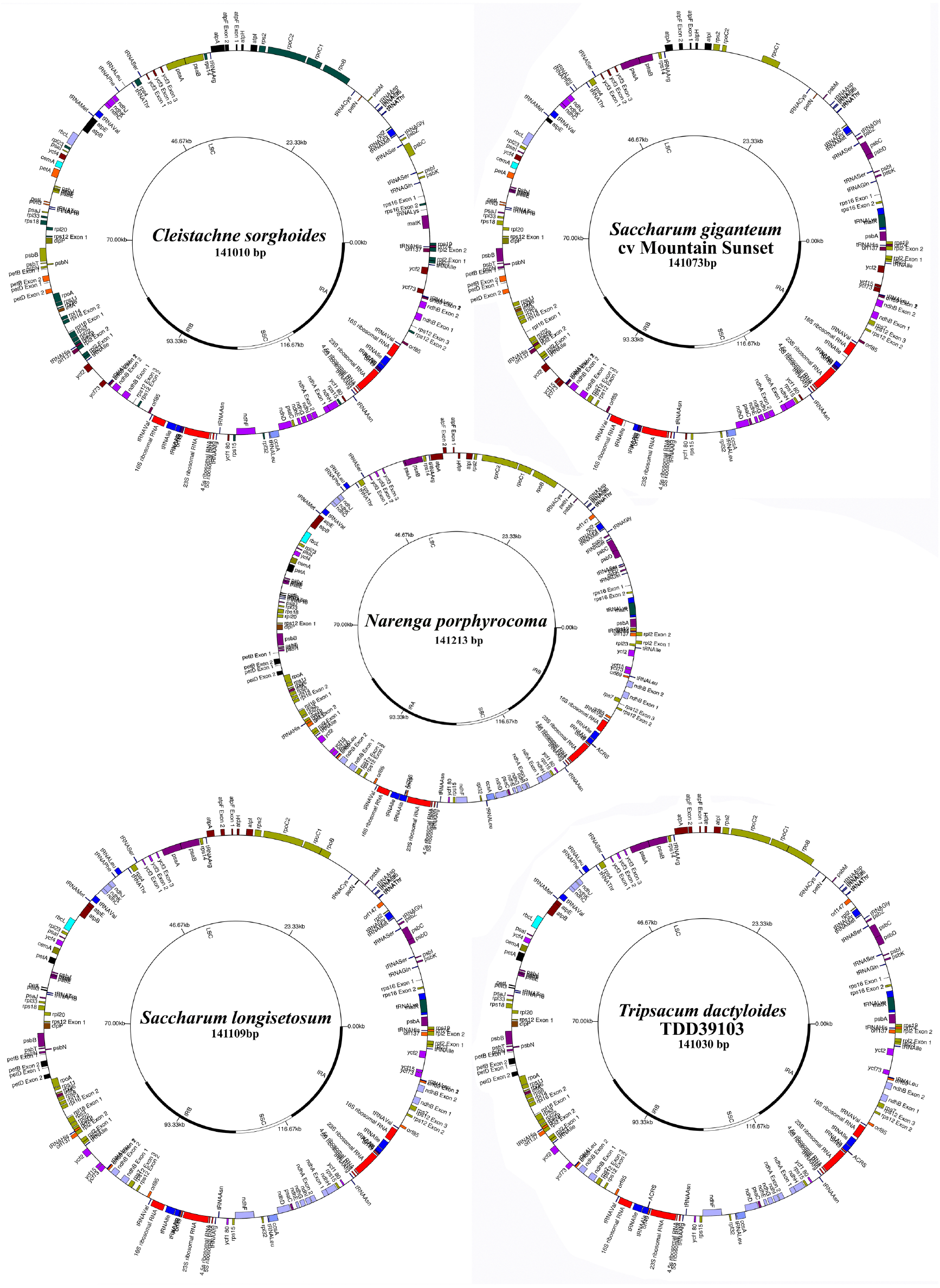
Schematic diagrams of five assembled chloroplasts. Schematic diagrams of the five main chloroplast genomes assembled in this study: *Cleistachne sorghoides*, *Saccharum giganteum* cv Mountain sunset, *Saccharum longisetosum, Narenga porphyrocoma* and *Tripsacum dactyloides* cv TDD39103. Sizes of each chloroplast genome are shown. Within the images, the inner track shows the location and sizes of the large single copy region (LSC), the first inverted repeat (IR_B_), the small single copy region (SSC) and the second inverted repeat (IR_A_). Chloroplast genes are shown on the outer track, with forward strand genes on the outside and reverse strand genes on the inside. Comparison images of the assembled chloroplast genomes with inverted SSC regions are shown in Supplementary Document 1.

As described previously, (Lloyd Evans et al. 2019) phylogenomic analysis of whole chloroplast genomes typically have good branch support (Figure 2) using a fine partitioning scheme, as presented in this paper. Overall, phylogenies are compatible with previous studies (Lloyd Evans et al. 2019, Estep et al. 2012). As expected, *Tripsacum dactyloides* is sister to *Zea mays* and *Zea luxurians*. *Cleistachne sorghoides* clusters within the Sorghum/Sarga complex, as an evolutionarily distal sister to *Sarga versicolor* and *Sarga timorense*. This grouping is sister to *Saccharum* and *Miscanthus*.

**Figure 2.**
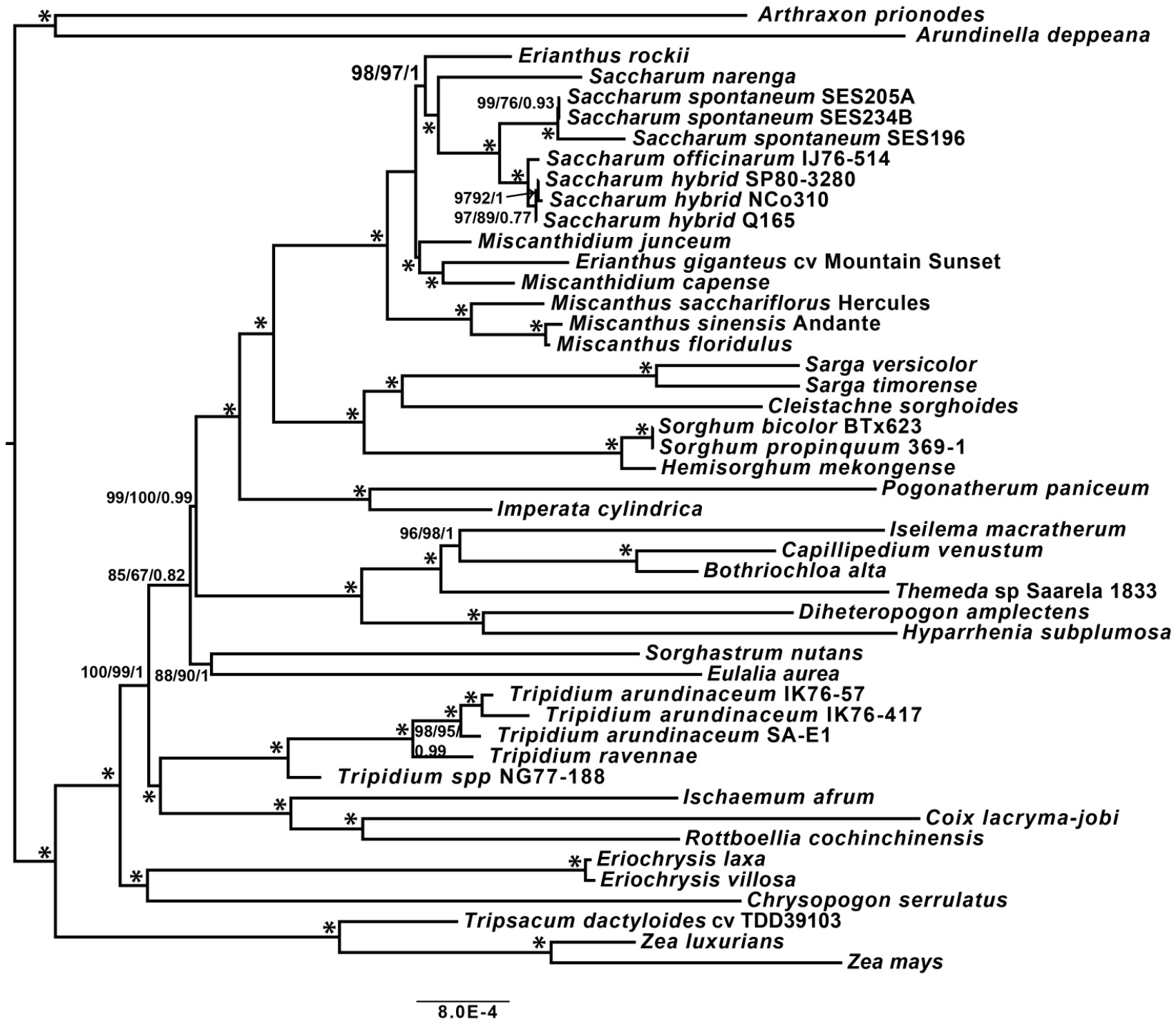
Whole chloroplast genome phylogeny. Whole chloroplast genome phylogeny centred on the Saccharinae, Sorghinae and core Andropogoneae. Numbers at nodes represent sh-aLRT single branch tests, non-parametric bootstrap tests and Bayesian inference. A* represents branches with 100% support across all tests.

*Erianthus giganteus* clusters within *Miscanthidium*. Interestingly, *Erianthus rockii* is an outgroup to both *Narenga* and *Saccharum*. As expected, *Saccharum spontaneum* is sister to *Saccharum officinarum* and modern hybrid cultivars (based on *Saccharum cultum* (Lloyd Evans and Joshi 2016)).

Within the extended ITS phylogeny (1200bp), the topology is essentially identical to previous work (Snyman et al. 2018). As described previously, *Sorghum* and *Sarga* split into distinct genera, with *Sorghum* sister to the core Andropogoneae and *Sarga* sister to *Miscanthus* and *Saccharum*. *Cleistachne sorghoides* remains as sister to the crown *Sarga* species whilst *Spodiopogon sibiricus* is sister to *Sorghum*. In the ITS phylogeny, *Erianthus rockii* and *Narenga porphyrocoma* form a clade that is sister to *Miscanthidium and Erianthus giganteus* forms an outgroup to *Miscanthidium capense* and *Miscanthidium junceum*. Branch support is generally good, but with lower support along the backbone of the tree.

## Discussion

Six novel chloroplast genomes from the Sorghinae and Sacchrinae were assembled from long read MinION ONT data. The relative ease and low cost of assembly ONT MinION sequencing (<$1800 for all the sequencing in this paper) makes the use of whole chloroplasts as an universal plant barcode feasible on a large scale for the first time. In parallel, full length ITS cassette sequences were amplified for the five species whose DNA was collected as well as the sugarcane cultivar SP80-3280.

The whole chloroplast and ITS assemblies were combined with public data for phylogenomic and phylogenetic analysis. For the whole chloroplasts assemblies a combined whole chloroplast alignment and phylogeny was possible. However, due to the relatively short ITS sequences currently available in the public domain, only a 2200bp alignment was feasible (even then many sequences had to be padded with Ns to fill in the termini). Overall, the phylogenetic analyses presented are compatible with previous results (Estep et al. 2012; Welker et al. 2015; Snyman et al. 2018; Lloyd Evans et al. 2019).

Addition of five extra species to the chloroplast phylogeny (Figure 2) has not altered the topology as compared with our previous analysis and the addition of the chloroplasts from six additional species has merely enriched the view on the Andropogoneae as a whole. Trivially, *Tripsacum dactyloides* emerges as sister to *Zea mays* and *Zea luxurians*. *Sarga timorense* is sister to *Sarga versicolor*, adding to the phylogeny of genus *Sarga*. *Cleistachne sorghoides* is sister to both *Sarga timorense* and *Sarga versicolor*, showing that this species is more closely related to *Sarga* rather than *Sorghum*. New World *Erianthus giganteus* (syn *Saccharum giganteum*) is placed within genus *Miscanthidium,* demonstrating that this species and genus is clearly not part of *Saccharum*. Interestingly, *Narenga porphyrocoma* emerges as sister to *Saccharum sensu stricto* and *Erianthus rockii* is sister to a clade formed by *Narenga* and *Saccharum*.

However, it should be noted that many species within the Andropogoneae have undergone reticulate (network) evolution where the chloroplast phylogeny can be highly divergent from the nuclear phylogeny. As a result, chloroplast derived phylogenies need to be compared with nuclear based phylogenies to reveal areas of correspondence and divergence.

The extended ITS phylogeny (2200 characters) is largely congruent with the previously published low copy number gene phyogenies (Lloyd Evans et al 2019, Estep et al. 2014) covering the Andropogoneae. As in the low copy number phylogenies, internal branches are well supported, whilst branches representing the backbone of the tree are more poorly supported, indicating rapid radiation during the evolution of the Andropogoneae. As a genus, *Microstegium* is clearly polyphyletic. Within the ITS phylogeny and the chloroplast phylogeny *Chleistachne sorghoides* emerges as an outgroup to *Sarga*. This would fit with the model that *Cleistachne sorghoides* arose from a genome duplication of an n=5 ancestor which as the common ancestor of *Sarga* with two chromosomes fusing to give n=9 as the base chromosome number.

As in previous chloroplast-based phylogenies, *Sorghum* and *Sarga* are sister lineages. However, in the ITS phylogeny *Sorghum* is sister to the core Andropogoneae, with *Spodiopogon sibiricus* sister to the Sorghum + Core Andropogoneae clade, whilst *Sarga* is sister to *Miscanthidium* and *Saccharum*. *Saccharum perrieri* and *Lasorhachis hildebrandii* are sister to the crown *Sorghums*, indicating that these are members of sorghum and should neither be part of *Saccharum* nor given their own genus. This, once again, reveals the confusion of nomenclature within both *Saccharum* and *Sorghum*.

The phylogeny supports the independence of *Tripidium* (Lloyd Evans et al. 2019) and demonstrates that, as a lineage, it is independent from *Germania*.

In the whole chloroplast phylogeny *Erianthus giganteus* is sister to *Miscanthidium capense* within the *Miscanthidium* clade. In the ITS phylogeny (Figure 3) *Erianthus giganteus* is sister to both *Miscanthidium*. *Erianthus rockii* is sister to *Narenga porphyrocoma*, with this clade being sister to *Miscanthidium*+*E. giganteus*. The low copy number gene locus phylogeny of Welker et al. (2015), which included several South American *Erianthus* species, (including *Erianthus giganteus*) placed *E. giganteus* as sister to a clade formed by *Narenga porphyrocoma B* and *Saccharum* as well as sister to a clade formed by *Narenga porphyrocoma A* and *Miscanthidium capense* (syn *Saccharum ecklonii*). This is a good indication that *Erianthus giganteus* is an hybird formed from a species ancestral to *Miscanthidium* and a species ancestral to *Saccharum* and *Narenga*. Our chloroplast phylogeny places *E. giganteus* within *Miscanthidium*, whilst the ITS phylogeny places *E. giganteus* as sister to *Miscanthidium*. It is obvious that *E. giganteus* is an hybrid, formed as a result of reticulate evolution, with part of its genome and its chloroplast originating from a species sister to *Saccharum*. The other part of the genome originates from a species sister to *Miscanthidium*. The hybrid nature of *E. giganteus* is different from that of *Miscanthidium*, suggesting that *E. giganteus* is distinct from *Miscanthidium* and should not be included in *Miscanthidium*. It is also separate from *Saccharum* and inclusion of *E. giganteus* (as *Saccharum longisetosum*) within *Saccharum* is erroneous. The most parsimonious solution is that *Erianthus* should be the preferred genus name for this species. As *E. giganteus* is the type species for *Erianthus*, this means that the other *Erianthus* species from the Americas should be removed from *Saccharum* and placed in their own genus, *Erianthus*.

**Figure 3.**
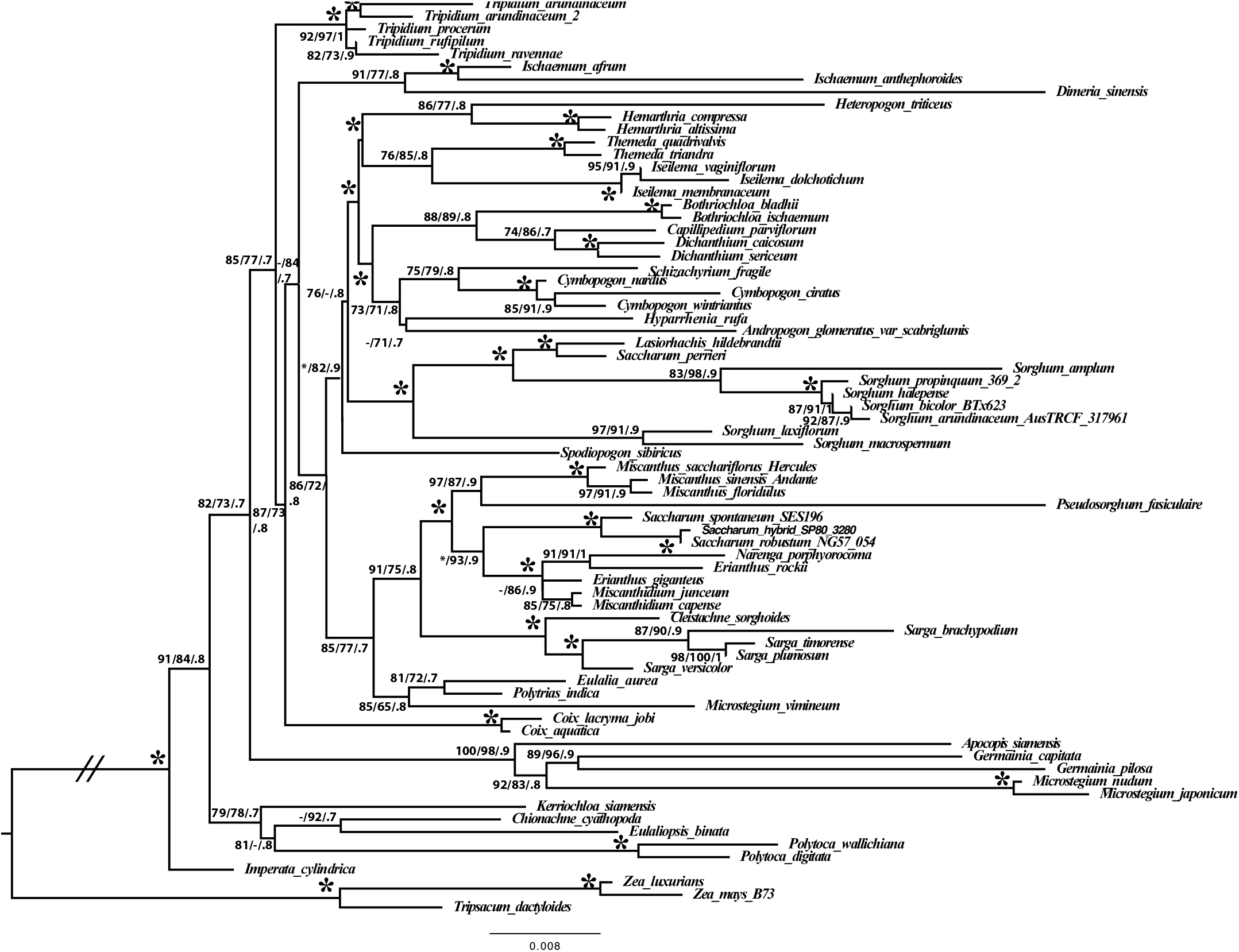
Extended ITS Phylogeny Phylogeny generated from an alignment of extended its regions (1200bp). Numbers at nodes represent sh-aLRT single branch tests/non-parametric bootstraps/Bayesian inference. A * represents 100% confidence in a branch, whilst – represent values below the threshold (80% for SH-alrt, 75% for bootstraps and 0.7 for Bayesian inference. Scale bar at the bottom represents the expected number of substitutions per site. The Tripsacinae were employed as an outgroup.

In our phylogenies, *Erianthus rockii* (syn *Erianthus longisetosum*) emerges as sister to both *Narenga* and *Saccharum* in the chloroplast phylogeny and is sister to *Narenga porphyrocoma* in the ITS phylogeny. The only other phylogenetic analysis to include *E. rockii* is the ITS-based phylogeny of Hodkinson et al. (2002). Though they only employed half the number of characters as compared with this study, and many branches collapsed into polytomies, *E. rockii* emerged as sister to *Miscanthus fuscus* and *Narenga porphyrocoma* with this grouping sister to *Miscanthidium* and *Saccharum contortum*. No low copy number gene phylogeny has yet been performed for *E. rockii*, though the data in this study suggests that it is an hybrid with a different and separate reticulate origin from *Narenga porphyorcoma* and *Erianthus giganteus*. Thus the species is not a member of genus *Saccharum*, though its precise taxonomic placement will have to wait for further work on low copy number genes or other genomic regions.

*Naenga porphyrocoma* emerges as sister to *Saccharum* in the chloroplast phylogeny, though in the ITS analysis it is sister to *Erianthus rockii* and *Erianthus giganteus*, with the entire grouping being sister to *Miscanthidium*. This agrees well with the low copy number gene phylogeny of Welker et al. (2015) and gives strong support to *Narenga porphyorcoma* being a hybrid of an immediate ancestor of *Saccharum* with a member of genus that is sister to *Miscanthidium*. Thus *Narenga porphyorcoma* has a separate evolutionary history from the other species analyzed in this study. It is not part of genus *Saccharum*, neither is it part of *Miscanthidium* or *Erianthus*.

Combining all the information above with the analyses of Welker et al. (2015) leads to a model of reticulate evolution in the species of interest that is shown in Figure 4. Starting from an n=5 *Sarga* genome it is possible, by combining ITS, chloroplast and low copy number genomic data, to trace the reticulate evolutionary origins of the saccharinae (species that are within the 3.5 million year window of divergence from *Saccharum* where hybridization in the wild is possible (Lloyd Evans and Joshi 2016)). The first to emerge is an A genome, of n=5 which is shared between *Mischantidium*, *Erianthus*, *E. rockii* and *Saccharum narenga*. A genome duplication, hybridization or retention event leads to the common (n=10) ancestor of *Miscanthus* and *Saccharum*. It should be noted that though *Sarga* has a base chromosome number of 5, this may have arisen through a whole genome hybridization (going from x=10 to x=5) resulting in larger chromosomes (Price et al. 2005). The *Miscanthus* lineage undergoes a hybridization and chromosome fusion event leading to the x=19 base chromosome number in *Miscanthus* (Ma et al. 2012).

**Figure 4.**
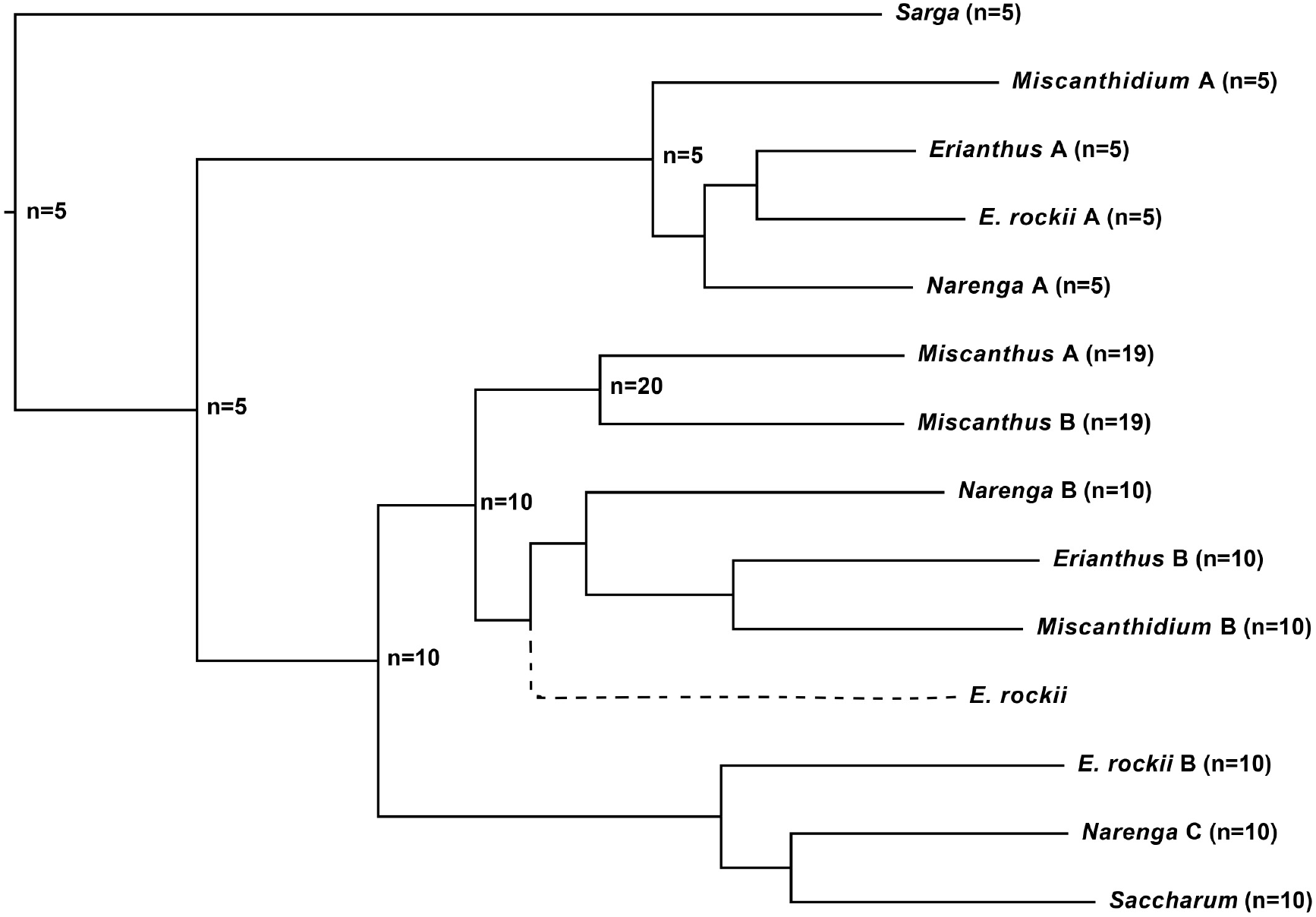
Model for the genome evolution of the Saccharinae. Schematic demonstrating the probably evolution of different genomes within the Saccharinae. The evolutionary series begins with Sarga with a base chromosome number of 5. Different lineages and proven genome ancestries are shown. A possible additional genome representation with x=10 sister to Miscanthus is shown with a dashed line as it cannot be proven with existing data.

The x=10 ancestral *Saccharum* lineage splits into two families, both with x=10. The first of these is the B genome of *Erianthus*, *Miscanthidium* and *Narenga*. The second lineage, also x=10 leads to the chromosomal lineages of *E. rockii*, *Narenga* and *Saccharum* (the B genome of *E. rockii* and the C genome of *Narenga*). Two chromosomal fusions in *Saccharum spontaneum* yields the x=8 base chromosomal number of this species. *Saccharum officinarum* retains its x=10 base chromosomal number, but undergoes multiple rounds of polyploidism.

Both *Miscanthidium* and *Erianthus* arose from hybridizations between an x=5 A genome and an x=10 B genome (though from different lineages) yielding a base chromosome number of x=15. They are closely related, but evolutionarily distinct. Thus, though looking at whole chloroplast genome, ITS and low copy number gene phylogenies independently might suggest that *Miscanthidium* and *Erianthus* should be combined in to a single genus, the separate and distinct origins of these genera indicates that they should be treated separately. In addition, *Miscanthidium* is an African genus whilst *Erianthus* is a genus from the Americas. *Miscanthidium* has three distinct ancestral genomes, x=5, x=10 and x=10 though rearrangement and chromosomal loss yielded a final base chromosomal number of 15. *Erianthus rockii* has the A genome of *Narenga*, and has a B genome that is sister to the C genome of *Narenga* (as well as the genome of *Saccharum*). However, without low copy number phylogenetics it is not known if *E. rockii* has a additional genome genome component equivalent to the B genome of *Narenga*. Clearly, *Narenga* is separate from *Saccharum* and the combination of an x=5 A genome and an x=10 B genome that is evolutionarily separate from other genera also makes *E. rockii* potentially a separate genus, though the precise nature of its reticulate evolution requires additional and separate analysis.

Surprisingly, of all the genera studied *Saccharum* is the only one not to demonstrate any signals of reticulate evolution. This gives the lie to *Sacharum* having arisen as the result of introgression from multiple species, as has been the predominant theory of Saccharum’s origins for over half a century (Mukherjee 1957). The data presented in this paper demonstrate that though *Erianthus*, *E. rockii* and *Narenga* all lie within the saccharinae (as a possible interbreeding group of genera related to *Saccharum*) they are not part of genus *Saccharum* itself.

These findings have considerable implications for phylogenetic analyses performed from chloroplast analyses alone (particularly in the grasses), as reticulate evolution can result a skewed view of grass origins if only using plastomes or plastome fragments for phylogenetic analyses. Moreover, as ITS and low copy number genes also provide different evolutionary signals combining ITS and chloroplast genes or regions can significantly muddy the waters. Multiple signals from different sources need to be analyzed separately and interpreted together to determine a more accurate phylogeny of the Andropogoneae. The most meaningful case in point is the relationship of *Sorghum* and *Sarga*. Chloroplast data consistently places these as closely related sister genera. However both low copy number nuclear genes and extended ITS sequences place *Sorghum* as sister to the core Andropogoneae and *Sarga* as sister to the Saccharinae, extending the evolutionary distance between them from 7.6 to 12.4 million years (Hawkins et al. 2015, Lloyd Evans and Joshi 2016; Snyman et al. 2018).

## Conclusion

The complexity of phylogenetic relationships within the Andropogoneae, particularly those species closest to sugarcane, is again revealed. ITS analysis confirms the results from low copy number gene studies (Hawkins et al. 2015; Lloyd Evans et al. 2019). Genus *Sarga* is distinct from *Sorghum* and forms an outgroup to *Saccharum* and allies. Indeed, *Sarga* may well delimit the taxonomic extent of the Saccharinae. African *Miscanthidium* is sister to *Saccharum* and *Miscanthus* is sister to the clade formed by *Miscanthidium* and *Saccharum*. By combining chloroplast and ITS phylogenetics with previously-published low copy number gene phylogenetics, a new model of the origins of the Saccharinae (Figure 4) is proposed. It is also demonstrated that *Erianthus*, Miscanthidium and *Narenga* have independent evolutionary paths based on separate origins (though *E. longisetosum* is sister to *Narenga* in both phylogenies). This and their different base chromosome numbers (x=15 as opposed to x=10 for *Saccharum*) means that they should be excluded from genus *Saccharum*. However, the genera *Sarga*, *Miscanthus*, *Miscanthidium*, *Erianthus*, *Narenga* as well as *E. longisetosum* are monophyletic (not being introgressed into each other) and should be considered as being part of the Saccharinae subtribe.

The taxonomy of the Saccharinae (and the Andropogoneae in general) cannot be fully understood without understanding the reticulate evolution of the species involved. Indeed, hybridizaton seems to be a common factor in speciation with multiple independent genomes being involved in this process. A single molecular phylogenetic technique cannot reveal the full evolutionary history of many species and genera and this has led to the incorrect and inappropriate inclusion of too many species into genus *Saccharum*. This paper takes a long step towards the final circumscription both of genus *Saccharum* and the Saccharinae subtribe.

## Acknowledgements

We would like to thank Mr K-L Ma for collecting the *S. timorense* seeds and the *N. porphyrocoma* leaf material. We are grateful to Oxford Nanopore Technologies for support through their community access programme. We also thank the British Association of Sugar Technologists (BAST) for providing *Saccharum* hybrid cv SP80-3280 plant material.

## Author Contributions

DLlE conceived the study, performed the bioinformatics, analyzed the data and wrote the paper. BH and DLLE designed the capture primers. BH performed the DNA isolation and sequencing. All authors read and approved the final version of the paper.

## Data Availability

Alignments and reference phylogenies are available from Zenodo (DOI: 10.5281/zenodo.2596381). Assembled and annotated chloroplast genomes have been deposited in ENA under the project accession PRJEB31602. Sequenced and annotated ITS regions have been deposited in ENA under the project accession PRJEB31603. All computer code developed for this project is available from GitHub: https://github.com/gwydion1/bifo-scripts.git.

## Supplementary Materials

**Supplementary document 1.** Comparisons of the canonical and inverted SSC forms of the chloroplast genomes assembled for this study.

**Supplementary table 1**. List of accessions and sources for sequences employed in phylogenetic analyses.

The table gives a list of accessions, sources and references for all chloroplast and ITS regions employed in this study.

## References

Adati S (1958) Studies on the genus *Miscanthus* with special reference to the Japanese species suitable for breeding purposes as fodder crops. Bull. Fac. Agric. Mie Univ. 17:1–112

Álvarez I, Wendel JF (2003) Ribosomal ITS sequences and plant phylogenetic inference. Molecular Phylogenetics and Evolution, 29:417–434.

Andersson NJ (1855) Öfversigt af Förhandlingar: Kongl. Svenska Vetenskaps-Akademien 12:163–164

Bentham G (1882) In: Hooker’s Icones Plantarum 14:t. 1379.

Besnard G, Christin PA, Malé PJG, Coissac E, Ralimanana H, Vorontsova MS (2013) Phylogenomics and taxonomy of *Lecomtelleae* (Poaceae), an isolated panicoid lineage from Madagascar. Annals of Botany, 112:1057–1066.

Burner DM (1991) Cytogenetic analyses of sugarcane relatives (Andropogoneae: Saccharinae). Euphytica, 54:125–133.

Celarier RP (1958) Cytotaxonomy of the Andropogoneae: III. Subtribe Sorgheae, genus Sorghum. Cytologia 23:395–418

Chen X, Zhou J, Cui Y, Wang Y, Duan B, Yao H (2018) Identification of *Ligularia* herbs using the complete chloroplast genome as a super-barcode. Frontiers in Pharmacology. 9:695.

Clayton WD, Vorontsova MS, Harman KT, Williamson H (2002 onwards) World Grass Species: Synonymy. http://www.kew.org/data/grasses-syn.html. Accessed 07 March 2019.

Connor HE (2004) Flora of New Zealand — Gramineae supplement I: *Danthonioideae*. New Zealand Journal of Botany. 2004. 42:771–795.

Darriba D, Taboada GL, Doallo R, Posada D (2012) jModelTest 2: more models, new heuristics and parallel computing. Nature Methods; 9:772.

Engels B (2015). Amplify4. Available from: https://github.com/wrengels/Amplify4 (last accessed March 3 2020).

Estep MC, McKain MR, Diaz DV, Zhong J, Hodge JG, Hodkinson TR, Layton DJ, Malcomber ST, Pasquet R, Kellogg EA (2014) Allopolyploidy, diversification, and the Miocene grassland expansion. Proceedings of the National Academy of Sciences, 111:15149–15154.

Folk RA, Mandel JR, Freudenstein JV (2017) Ancestral gene flow and parallel organellar genome capture result in extreme phylogenomic discord in a lineage of angiosperms. Systematic Biology. 66:320–337.

Gu MH, Ma HT, Liang GH (1984) Karyotype analysis of seven species in the genus *Sorghum*. J. Hered. 75:196–202

Ha S, Moore PH, Heinz D, Kato S, Ohmido N, Fukui K (1999) Quantitative chromosome map of the polyploid *Saccharum spontaneum* by multicolor fluorescence in situ hybridization and imaging methods. Plant Mol. Biol. 39:1165–1173

Hawkins JS, Ramachandran D, Henderson A, Freeman J, Carlise M, Harris A, Willison-Headley Z (2015) Phylogenetic reconstruction using four low-copy nuclear loci strongly supports a polyphyletic origin of the genus Sorghum. Annals of Botany, 116:291–299.

Hinsinger DD, Gaudeul M, Couloux A, Bousquet J, Frascaria-Lacoste N (2014) The phylogeography of Eurasian *Fraxinus* species reveals ancient transcontinental reticulation. Molecular Phylogenetics and Evolution. 77:223–237.

Hodkinson TR, Chase MW, Lledó DM, Salamin N, Renvoize SA (2002) Phylogenetics of Miscanthus, Saccharum and related genera (Saccharinae, Andropogoneae, Poaceae) based on DNA sequences from ITS nuclear ribosomal DNA and plastid trnL intron and trnL-F intergenic spacers. Journal of Plant Research, 115:381–392.

Jensen KB, Highnight K, Wipff KJ (1989) IOPB chromosome data 1. Int. Organ. Pl. Biosyst. Newslett. (Zurich) 13:20–21.

Kellogg EA, Appels R, Mason-Gamer RJ. (1996) When genes tell different stories: the diploid genera of Triticeae (Gramineae). Systematic Botany. 21:321–347.

Kellogg EA. (2013) Phylogenetic Relationships of Saccharinae and Sorghinae. In: Paterson AH, editor. Genomics of the Saccharinae. New York: Springer. pp. 3–21.

Keng YL (1939) The gross morphology of Andropogoneae (from the standpoint of taxonomy). Sinensia, 10:273–343.

Krawczyk K, Nobis M, Myszczyński K, Klichowska E, Sawicki J (2018) Plastid super-barcodes as a tool for species discrimination in feather grasses (Poaceae: Stipa). Scientific reports, 8:1924.

Li HW, Shang KC, Hsiao YY, Yang PC (1959) Cytological studies of sugarcane and its relatives. Cytologia, 24:220–236

Linnaeus C (1753) Species Plantarum. Imprensis Laurentii Salvii. Holmiae.

Liu K, Raghavan S, Nelesen S, Linder CR, Warnow T. 2009. Rapid and accurate large-scale coestimation of sequence alignments and phylogenetic trees. Science. 324:1561–1564.

Liu X, Fang F, Zhang R, Song H, Yang R, Gao Y, Ou H, Lei J, Luo T, Duan W, Zhang G (2012) Identification of progenies from sugarcane× Narenga porphyrocoma (Hance) Bor. by SSR marker. Southwest China Journal of Agricultural Sciences, 25:38–43.

Lloyd Evans D, Joshi SV (2016) Complete chloroplast genomes of *Saccharum spontaneum*, *Saccharum officinarum* and *Miscanthus floridulus* (Panicoideae: Andropogoneae) reveal the plastid view on sugarcane origins. Systematics and Biodiversity. 14:548–571.

Lloyd Evans D, Joshi SV, Wang J (2019) Whole chloroplast genome and gene locus phylogenies reveal the taxonomic placement and relationship of *Tripidium* (Panicoideae: Andropogoneae) to sugarcane. BMC Evolutionary Biology 19:33.

Lloyd Evans D (2020) Data from: Complete Chloroplast Genomes of *Saccharum giganteum*, *Saccharum longisetosum*, *Cleistachne sorghoides, Sarga timorense* and *Tripsacum dactyloides*. Placement within Saccharum and Comparisons with their ITS phylogeny. Zenodo. DOI: 10.5281/zenodo.2596381

Löve A (1976) IOPB Chromosome Number Reports LIV. Taxon. 25:631–649.

Löytynoja A, Vilella AJ, Goldman N (2012) Accurate extension of multiple sequence alignments using a phylogeny-aware graph algorithm. Bioinformatics. 28:1684–91.

Ma XF, Jensen E, Alexandrov N, Troukhan M, Zhang L, Thomas-Jones S, Farrar K, Clifton-Brown J, Donnison I, Swaller T, Flavell R (2012) High resolution genetic mapping by genome sequencing reveals genome duplication and tetraploid genetic structure of the diploid *Miscanthus sinensis*. PloS one, 7:p.e33821.

Maier RM, Neckermann K, Igloi GL, Kössel H (1995) Complete sequence of the maize chloroplast genome: gene content, hotspots of divergence and fine tuning of genetic information by transcript editing. Journal of Molecular Biology, 251:614–628.

Mukherjee SK (1957) Origin and distribution of Saccharum. Botanical Gazette, 119L55–61.

Narayanaswami V. In: Flora of Assam 5:461

Nguyen L-T, Schmidt HA, von Haeseler A, Minh QA (2015) IQ-TREE: A Fast and Effective Stochastic Algorithm for Estimating Maximum-Likelihood Phylogenies, Mol Biol and Evol. 32:268–274. doi:https://doi.org/10.1093/molbev/msu300.

Nylander JA, Wilgenbusch JC, Warren DL. Swofford DL (2008) AWTY (are we there yet?): a system for graphical exploration of MCMC convergence in Bayesian phylogenetics. Bioinformatics. 24:581–83.

Palisot de Beauvos AMFJ (1812) Essai d’une Nouvelle Agrostographie. Imprimerie de Fain. Paris.

Persoon CH. (1805) Synopsis Plantarum. Carol. Frid. Cramerum. Paris.

Piperidis G, Christopher MJ, Carroll BJ, Berding N, D’Hont A (2000). Molecular contribution to selection of intergeneric hybrids between sugarcane and the wild species *Erianthus arundinaceus*. Genome, 43:1033–1037.

Price HJ, Dillon SL, Hodnett G, Rooney WL, Ross L, Johnston JS (2005) Genome evolution in the genus *Sorghum* (Poaceae). Annals of Botany, 95:219–227.

Rambaut A, Suchard MA, Xie D, Drummond AJ (2009) Tracer, Version 1.5. http://tree.bio.ed.ac.uk/software/tracer/. Accessed 07 October 2017.

Ronquist F, Huelsenbeck JP (2003) MrBayes 3: Bayesian phylogenetic inference under mixed models. Bioinformatics. 19:1572–4.

Shepard AR, Rae JL. 1997. Magnetic bead capture of cDNAs from double-stranded plasmid cDNA libraries. Nucleic Acids Research, 25:3478–3480.

Snyman SJ, Komape D, Khanyi H, Van Den Berg J, Cilliers D, Lloyd Evans D, Barnard S, Siebert SJ (2018) Assessing the likelihood of gene flow from sugarcane (*Saccharum* hybrids) to wild relatives in South Africa. Frontiers in Bioengineering and Biotechnology, 6:72.

Stamakis A. 2006. RAxML-VI-HPC: Maximum likelihood-based phylogenetic analyses with thousands of taxa and mixed models. Bioinformatics. 22:2688–2690.

Sreenivasan TV, Sreenivasan J (1984) Cytology of Saccharum complex from New Guinea, Indonesia and India. Caryologia 37:351–357

Strydom A, Spies JJ, Van Wyk SMC (2000) Poaceae: Chromosome studies on African plants. 14. Panicoideae. Bothalia 39:201–214

Sukumaran J, Holder MT (2010) DendroPy: A Python library for phylogenetic computing. Bioinformatics. 26:1569–1571.

Walter T (1788) Flora Caroliniana, secundum. J. Fraser. London.

Wang W, Lanfear R. 2019. Long-reads reveal that the chloroplast genome exists in two distinct versions in most plants. Genome Biology and Evolution. 11:3372–3381.

Welker CA, Souza-Chies TT, Longhi-Wagner HM, Peichoto MC, McKain MR, Kellogg EA (2015) Phylogenetic analysis of *Saccharum* sl (Poaceae; Andropogoneae), with emphasis on the circumscription of the South American species. American Journal of Botany, 10:248–263.

Ye J, Coulouris G, Zaretskaya I, Cutcutache I, Rozen S, Madden TL (2012). Primer-BLAST: a tool to design target-specific primers for polymerase chain reaction. BMC Bioinformatics. 13:134.

Záveská E, Fér T, Šída O, Marhold K, Leong-Škorničková J (2016) Hybridization among distantly related species: examples from the polyploid genus *Curcuma* (Zingiberaceae). Molecular Phylogenetics and Evolution. 2016:100:303–321.

Zeng L, Zhang Q, Sun R, Kong H, Zhang N, Ma H (2014) Resolution of deep angiosperm phylogeny using conserved nuclear genes and estimates of early divergence times. Nature Communications 5:4956.

Zhu, Q (2014) BeforePhylo.pl version 0.9.0 available from: https://github.com/qiyunzhu/BeforePhylo. Accessed February 14 2019.

