## Supplementart Document 1 for "Complete Chloroplast Genomes of *Saccharum giganteum*, *Saccharum longisetosum*, *Cleistachne sorghoides, Sarga timorense, Narenga porphyrocoma* and *Tripsacum dactyloides*. Comparisons with ITS phylogeny and Placement within *Saccharum*"

### Supplementary Document 1

Chloroplast genomes assembled in this study from ONT MinION data

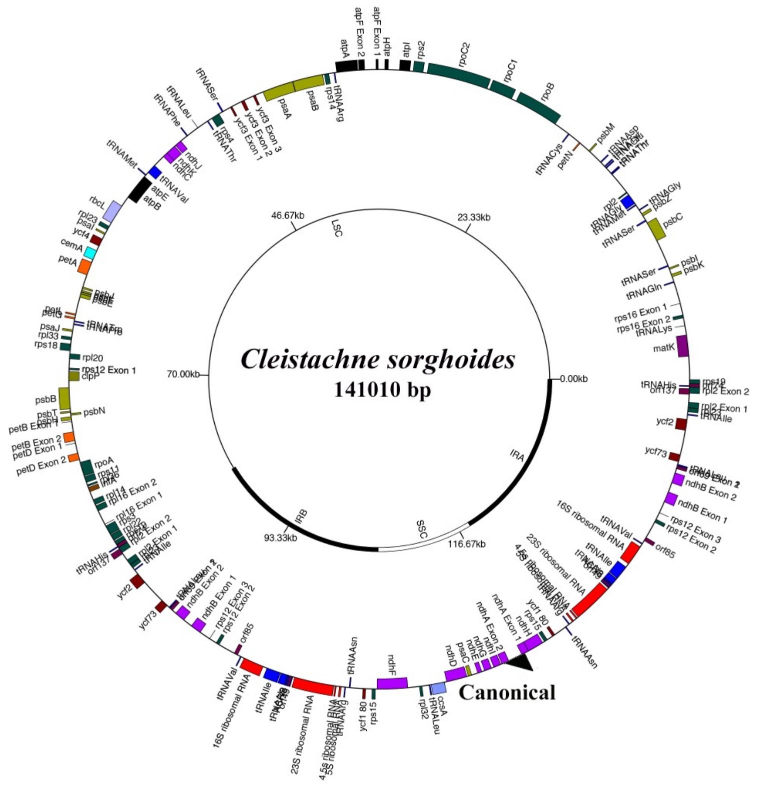


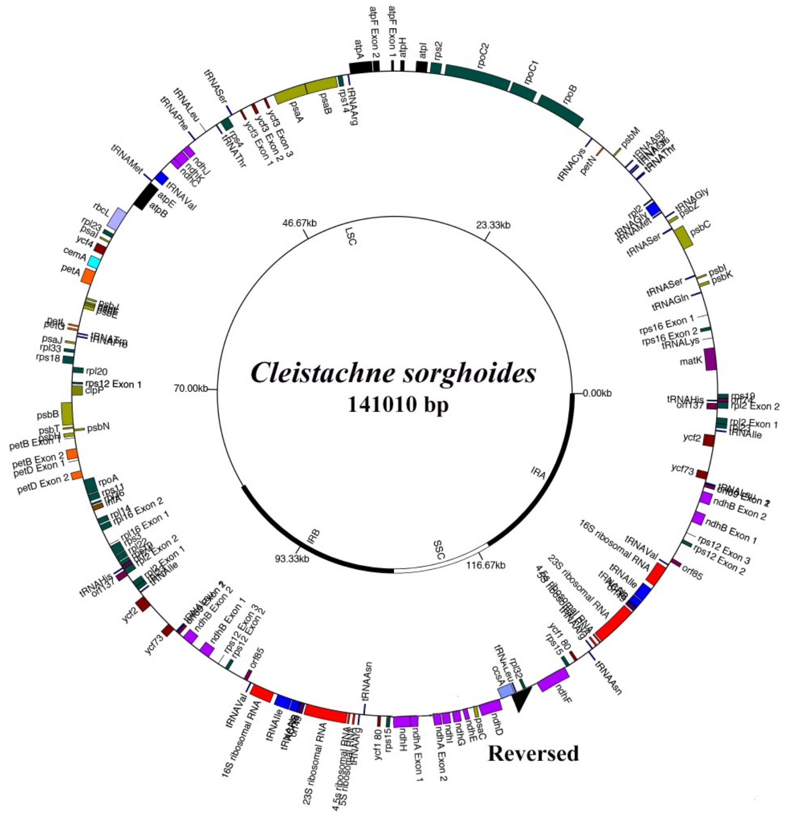


The two isoforms of the *Chleistachne sorghoides* chloroplast genome, with the canonical form (SSC in forward orientation) and reverse (SSC in reverse orientation) being present at a ratio of 51.4:48.6.

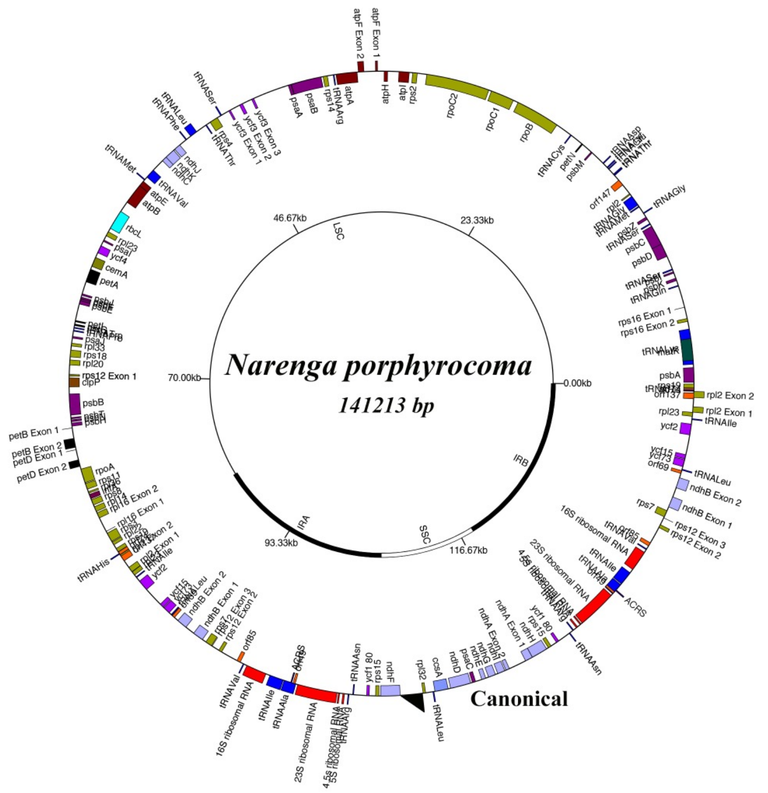


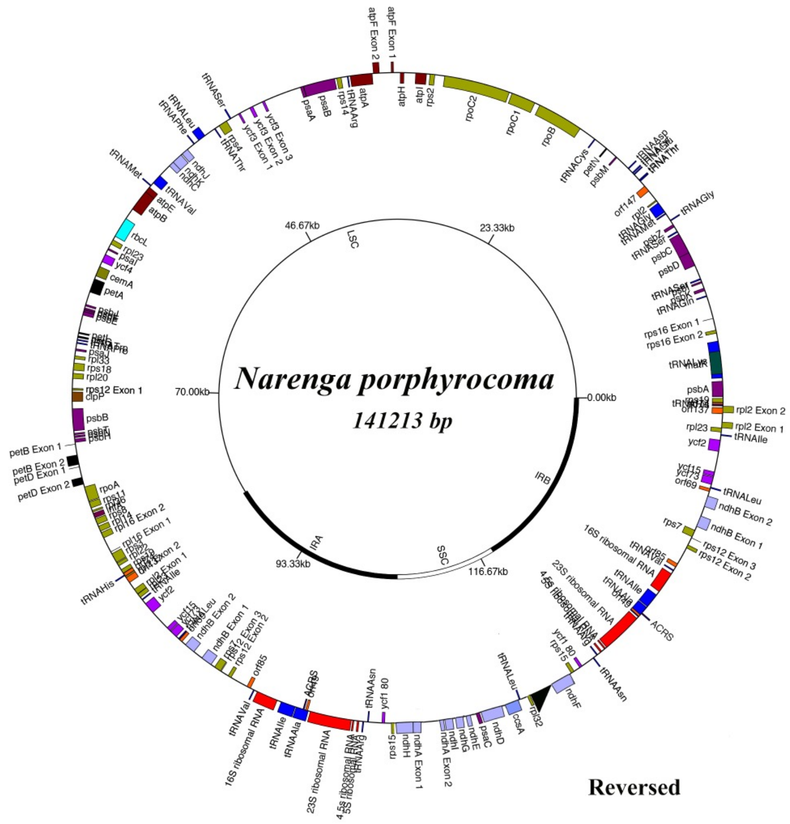


The two isoforms of the *Narenga porphyrocoma* chloroplast genome, with the canonical form (SSC in forward orientation) and reverse (SSC in reverse orientation) being present at a ratio of 47.2:52.8.

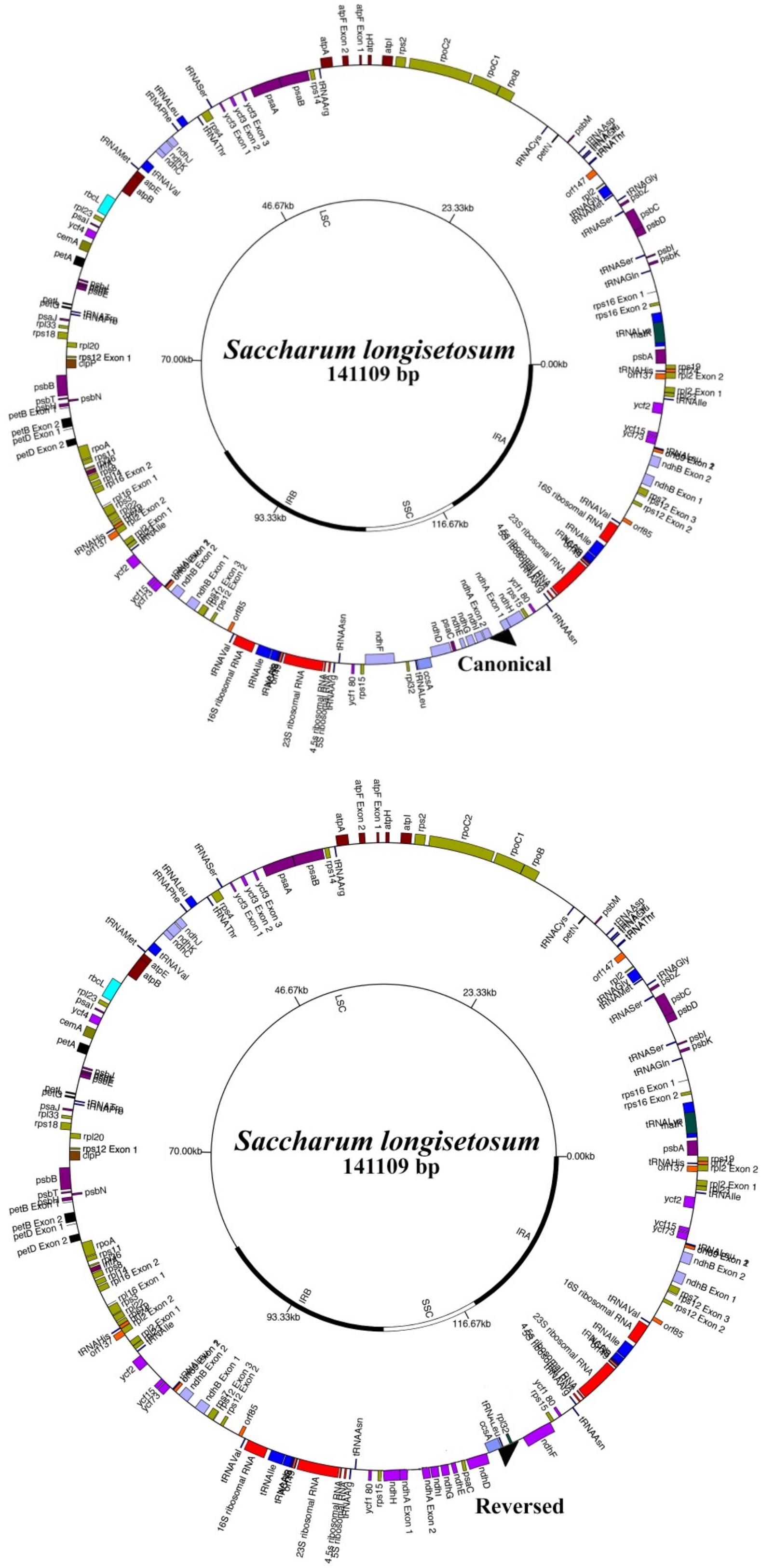


The two isoforms of the *Saccharum longisetosum (Erianthus rockii)* chloroplast genome, with the canonical form (SSC in forward orientation) and reverse (SSC in reverse orientation) being present at a ratio of 53.1:46.9.

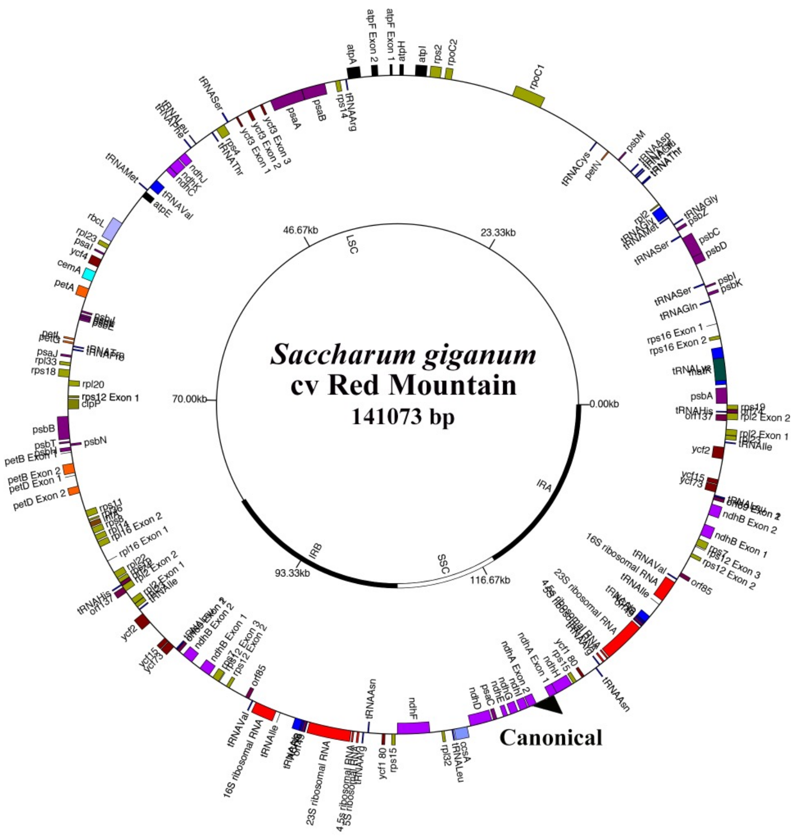


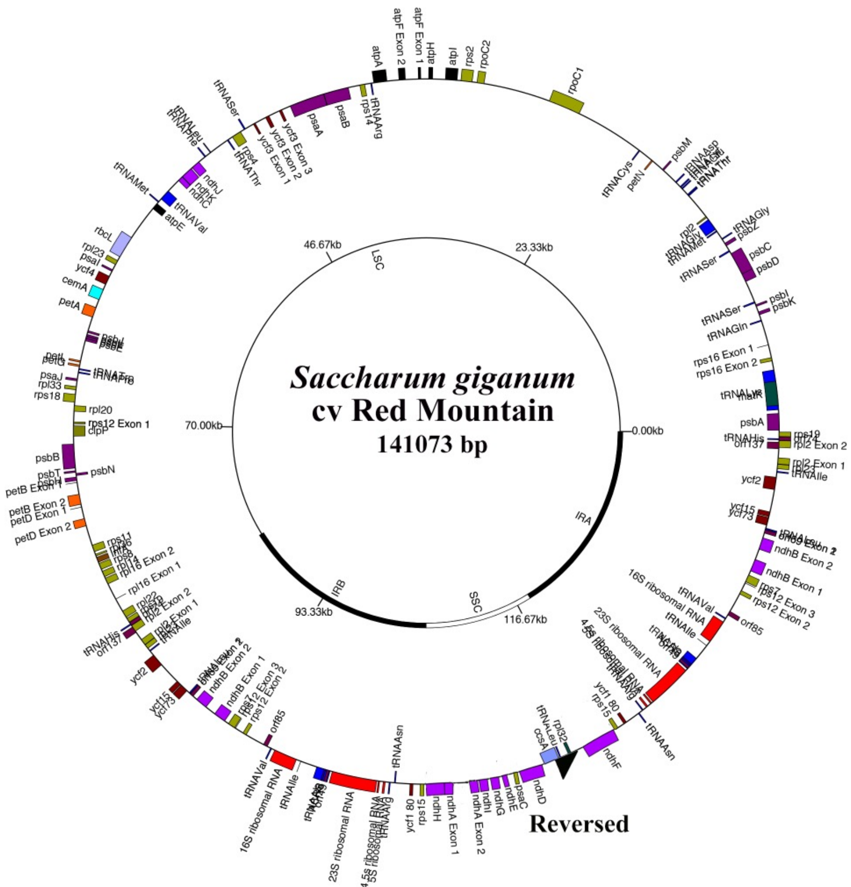


The two isoforms of the *Saccharum giganteum (Erianthus giganteus)* chloroplast genome, with the canonical form (SSC in forward orientation) and reverse (SSC in reverse orientation) being present at a ratio of 53.3:46.7.

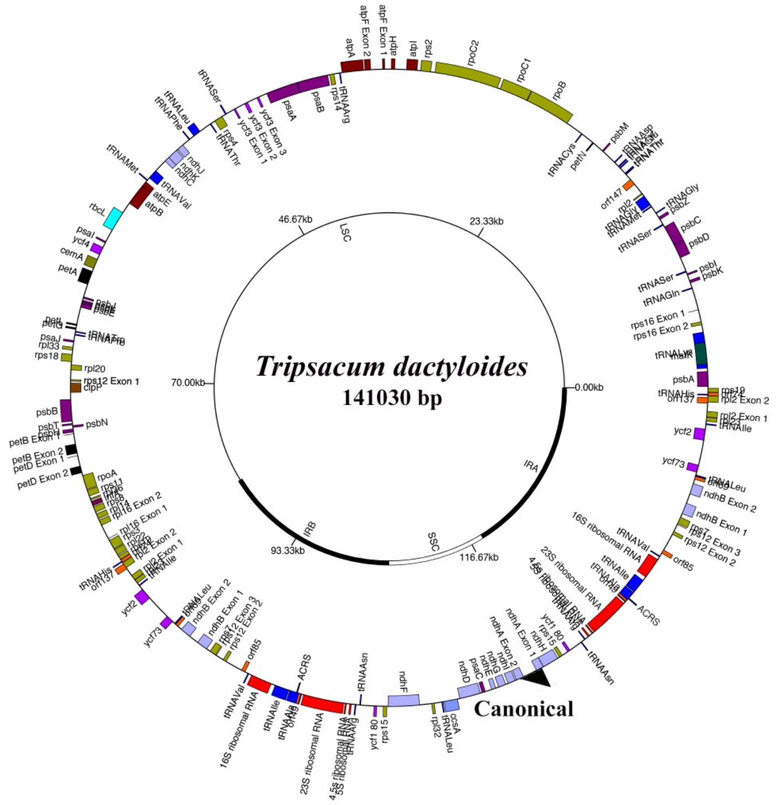


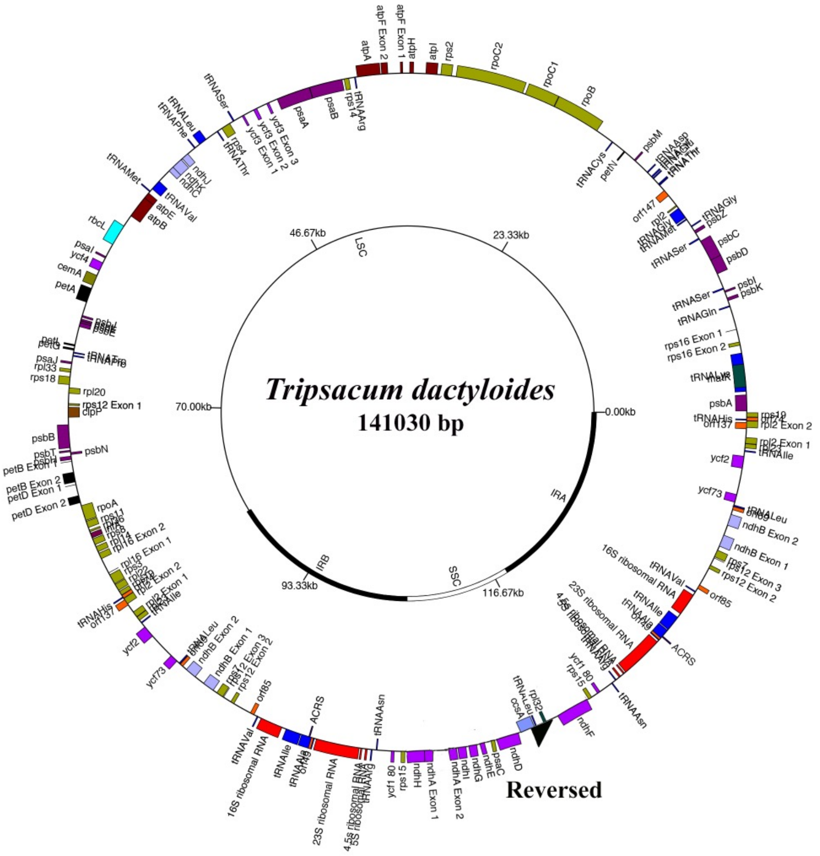


The two isoforms of the *Tripsacum dactyloides* chloroplast genome, with the canonical form (SSC in forward orientation) and reverse (SSC in reverse orientation) being present at a ratio of 50.4:49.6.

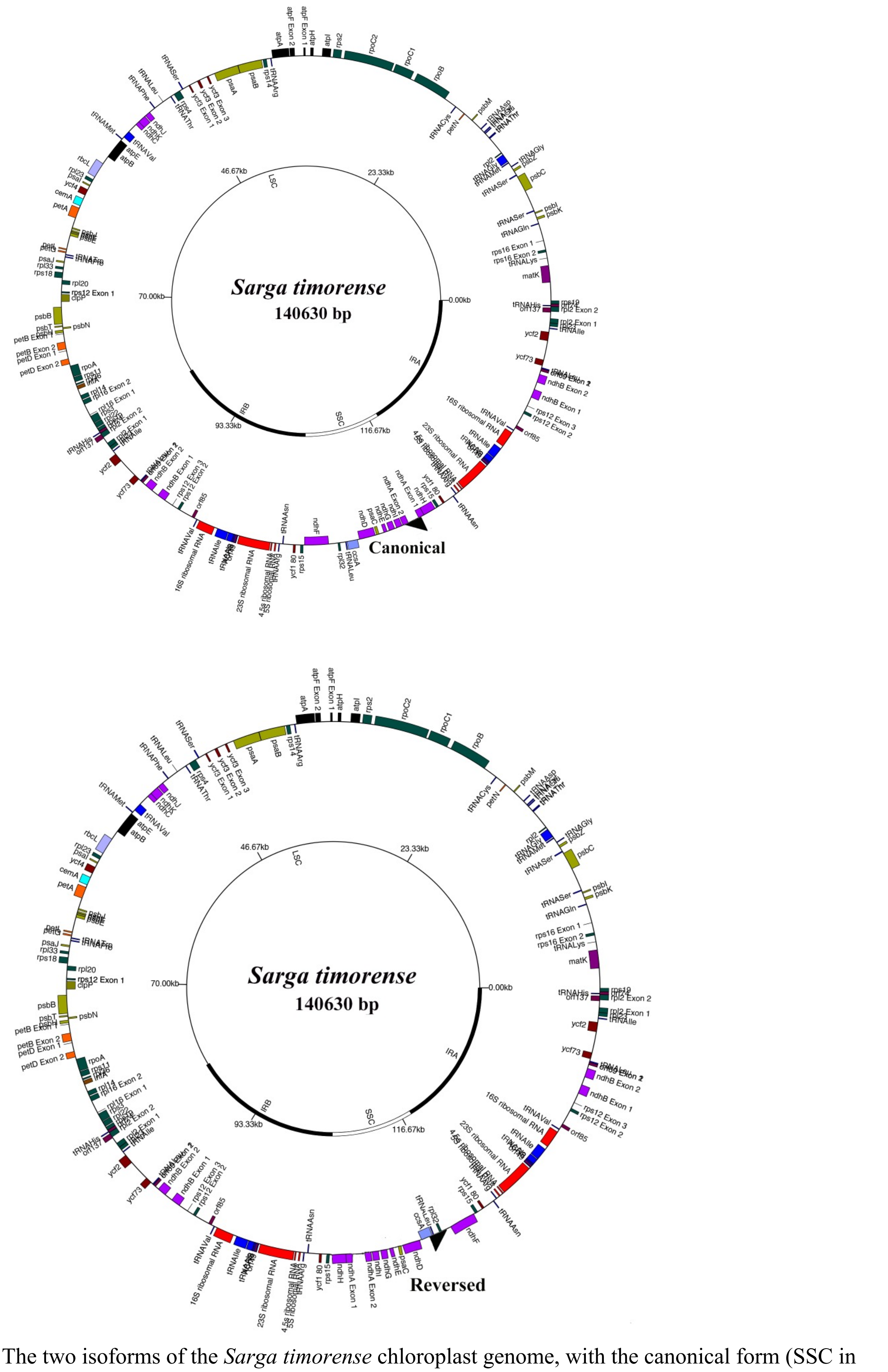


The two isoforms of the *Sarga timorense* chloroplast genome, with the canonical form (SSC in forward orientation) and reverse (SSC in reverse orientation) being present at a ratio of 51.1:48.9.
