## Supplementary Table 1 for "Complete Chloroplast Genomes of *Saccharum giganteum*, *Saccharum longisetosum*, *Cleistachne sorghoides, Sarga timorense, Narenga porphyrocoma* and *Tripsacum dactyloides*. Comparisons with ITS phylogeny and Placement within *Saccharum*"

| Species | Voucher or Accession | GenBank Accession | Reference |
|---|---|---|---|
| **Chloroplast Data** | | | |
| *Arundinella deppeana Nees ex Steud.* | XAL:Clark et al. 1680 | KU291490.1 | Burke et al. 2016 |
| *Arthraxon prionodes (Steud.) Dandy* | PI<ITA>:659331 | KU291471.1 | Burke et al. 2016 |
| Chrysopogon serrulatus Trin. | | KU961864.1 | Welker et al. 2016 |
| *Eriochrysis laxa* Swallen | | KU961863.1 | Welker et al. 2016 |
| *Eriochrysis villosa* Swallen | | KU961860.1 | Welker et al. 2016 |
| *Saccharum spontaneum* L. | SES205A | LN896360.1 | Lloyd Evans and Joshi 2016 |
| *Saccharum spontaneum* L. | SES234B | LN849912.1 | Lloyd Evans, D. (submitter) |
| *Saccharum spontaneum* L. | SES196 | PRJEB24335 | Lloyd Evans et al. 2019 |
| *Saccharum hybrid cultivar hort. Ex RM Grey* | SP80-3280 | AE009947.2 | Calsa Jr et al. 2004 |
| *Saccharum hybrid cultivar hort. Ex RM Grey* | Q165 | LN896359.1 | Lloyd Evans and Joshi 2016 |
| *Saccharum hybrid cultivar hort. Ex RM Grey* | | NCo310 | Asano et al. 2004 |
| *Saccharum officinarum* L. | IJ76-514 | LN849913.1 | Lloyd Evans and Joshi 2016 |
| *Miscanthidium junceum* Stapf (Stapf) | | LN869216.1 | Lloyd Evans, D. (submitter) |
| *Miscanthidium capense* (Nees) Stapf | Joshi 1 | PRJEB17861 | Lloyd Evans, D. (submitter) |
| *Miscanthus sacchariflorus (Maxim.) Benth. & Hook. f. ex Franch.* | cv Hercules | LN869218.1 | Lloyd Evans, D. (submitter) |
| *Miscanthus sinensis* Andersson | cv Andante | LS398101 | Lloyd Evans, D. (submitter) |
| *Miscanthus floridulus (Labill.) Warb. Ex K Schum. & Lauterb.* | PI295762 | LN869215.1 | Lloyd Evans, D. (submitter) |
| *Sorghum bicolor* (L.) Moench | BTx623 | EF115542.1 | Saski et al. 2007 |
| *Imperata cylindrical* (L.) Raeusch. | DEK:Burke 21 | KU291466.1 | Burke et al. 2016 |
| *Pogonatherum paniceum* (Lam.) Hack. | | KU961859.1 | Welker et al. 2016 |
| *Eulalia aurea* (Bory) Knuth | PI<ITA>:12153 | KU291499.1 | Burke et al. 2016 |
| *Sorghastrum nutans* (L.) Nash | DEK:Wysocki s.n. | KU291482.1 | Burke et al. 2016 |
| *Hyparrhenia subplumosa* Stapf | PI<ITA>:12665 | KU291500.1 | Burke et al. 2016 |
| *Diheteropogon amplectens Nees (Clayton)* | var. catangensis voucher PI<ITA>:12585 | KU291497.1 | Burke et al. 2016 |
| *Themeda sp.* Forssk. | Saarela 1833 | KU291484.1 | Burke et al. 2016 |
| *Iseilema macratherum Domin* | PI<ITA>:257760 | KU291468.1 | Burke et al. 2016 |
| Capillipedium venustum (Thwaites) Bor | PI<ITA>:11713 | KU291493.1 | Burke et al. 2016 |
| Bothriochloa alta (Hitchc.) Henrard | DEK:Duvall s.n. | KU291492.1 | Burke et al. 2016 |
| *Tripidium arundinaceum* Lloyd Evans | IK76-57 | PRJEB17861 | Lloyd Evans et al. 2019 |
| *Tripidium arundinaceum* Lloyd Evans | IK76-417 | PRJEB17861 | Lloyd Evans et al. 2019 |
| *Tripidium arundinaceum* Lloyd Evans | SA-E1 | PRJEB17861 | Lloyd Evans et al. 2019 |

| | | | |
|---|---|---|---|
| *Tripidium ravennae* | | PRJEB17861 | Lloyd Evans et al. 2019 |
| Tripidium sp. | NG77-188 | PRJEB17861 | Lloyd Evans et al. 2019 |
| *Ischaemum afrum* (JF Gmel.) Dandy | | PI<ITA>:36492 4 | Burke et al. 2016 |
| *Rottboellia cochinchinensis* (Lour.) Clayton | ISC<USA-IA>:Clark et al. 1698 | KU291481.1 | Burke et al. 2016 |
| *Coix lacryma-jobi* L. | M. Duvall s.n. 26May 2006 (DEK) | FJ261955.1 | Leseberg and Duvall 2009 |
| *Dimeria ornithopoda* Trin. | | KY596130.1 | Arthan et al. 2017 |
| *Eulaliopsis binata* (Retz.) CE Hubb. | | KY596182.1 | Arthan et al. 2017 |
| *Heteropogon triticeus* (R. Br.) Stapf ex Craib | | KY596142.1 | Arthan et al. 2017 |
| *Andropogon distachyon* L. | | KY596170.1 | Arthan et al. 2017 |
| *Schizachyrium sanguineum* (Retz.) Alston | | KY596124.1 | Arthan et al. 2017 |
| *Hemisorghum mekongense* (A. Camus) CE Hubb. | | KY596132.1 | Arthan et al. 2017 |
| *Eremochloa ciliaris* (L.) Merr. | | KY596146.1 | Arthan et al. 2017 |
| *Mnesithea helferi* (Hook. f.) de Koning & Sosef | | KY596162.1 | Arthan et al. 2017 |
| Sorghum propinquum (Knuth) Hitch. | 369-1 | PRJEB17862 | Lloyd Evans, D. (submitter) |
| *Zea mays* L. | B73 | AY928077.1 | Schnable et al. 2009 |
| *Zea luxurians* (Durieu & Asch.) RM Bird | | KR873424.1 | Orton 2015 |
| *Sarga versicolor* | | LS398104 | Lloyd Evans, D. (submitter) |

### ITS Data

| | | | |
|---|---|---|---|
| *Miscanthus oligostachyus* | | HQ822027 | Jang,J.-H. and Yoo,K.-O. (submitters) |
| *Miscanthus floridulus US56-0022-03* | | assembled from SRR486154 | Snyman et al. 2018 |
| *Miscanthus sinensis Andante* | | | ibid |
| *Polytoca digitata* | | GQ870232.1 | Teerawatananon et al. 2011 |
| *Polytrias indica* | | | Snyman et al. 2018 |
| *Microstegium vimineum 2* | | assembled from: ERR2040772 | Snyman et al. 2018 |
| *Bothriochloa insculpta* | | | Snyman et al. 2018 |
| *Andropogon glomeratus var scabriglumus* | | MF964041.1 | Thornhill et al. 2017 |
| *Andropogon virginicus* | | | ibid |
| *Hyparrhenia rufa* | | GQ870187.1 | Teerawatananon et al. 2011 |
| *Schizachyrium sanguineum* | | DQ005070.1 | Skendzic et al. 2007 |
| *Sorghastrum nutans* | | DQ005080.1 | Skendzic et al. 2007 |
| *Cymbopogon flexuosus* | | Assembled from: SRR2970609 | |
| *Sorghum ×drummondii* | | Assembled from: SRR998968 | |
| *Sorghum arundinaceum 2* | | Assembled from: SRR999026 | |
| *Sorghum halepense* 2 | | Assembled from: SRR486216 | |
| *Sorghum propinquum* | 369-1 | Assembled from: SRR998982 | |
| *Germainia capitata* | | GQ870198.1 | Teerawatananon et al. 2011 |
| *Microstegium japonicum* | | KF163847.1 | Kim et al 2012 |
| *Microstegium nudum* | | EU489073.1 | Chen et al. 2009 |
| *Saccharum arundinaceum 2* | | JX156345.1 | Zeng,H., Wei,L. and Liu,X. (submitters) |

| | | | |
|---|---|---|---|
| *Imperata cylindrica* | | MH768200.1 | Li et al. 2018 |
| *Zea mays* | B73 | Extracted from genome reference | |
| *Sorghum bicolor* | BTx623 | Extracted from genome reference | |
| *Sorghum laxiflorum* | | GQ121741.1 | Ng'uni et al. 2010 |


Arthan W, McKain MR, Traiperm P, Welker CA, Teisher JK and Kellogg EA. 2017. Phylogenomics of Andropogoneae (Panicoideae: Poaceae) of Mainland Southeast Asia. Systematic Botany, 42:418-431.

Asano T, Tsudzuki T, Takahashi S, Shimada H, Kadowaki KI. 2004. Complete nucleotide sequence of the sugarcane (Saccharum officinarum) chloroplast genome: a comparative analysis of four monocot chloroplast genomes. DNA Research 11:93–99.

Burke, S.V., Wysocki, W.P., Zuloaga, F.O., Craine, J.M., Pires, J.C., Edger, P.P., Mayfield-Jones, D., Clark, L.G., Kelchner, S.A. and Duvall, M.R., 2016. Evolutionary relationships in Panicoid grasses based on plastome phylogenomics (Panicoideae; Poaceae). *BMC Plant Biology*, *16*(1), p.140.

Calsa Jr T, Carraro DM, Benatti MR, Barbosa AC, Kitajima JP, Carrer H. 2004. Structural features and transcript-editing analysis of sugarcane (Saccharum officinarum L.) chloroplast genome. Current Genetics 46:366–373.

Chen CH, Veldkamp JF, Kuoh CS, Tsai CC, Chiang YC. 2009. Segregation of Leptatherum from Microstegium (Andropogoneae, Poaceae) confirmed by internal transcribed spacer DNA sequences. *Blumea-Biodiversity, Evolution and Biogeography of Plants*, *54*:175-180.

Kim CS, Yeon JS, Han YW, Lee IY, 2012. Plant DNA barcode and the case study of Korean Panicoideae, Poaceae. *한국잡초학회 별책 (학술대회 초록집)*, *32*:186-187.

Lloyd Evans D, Joshi SV, Wang J (2019) Whole chloroplast genome and gene locus phylogenies reveal the taxonomic placement and relationship of *Tripidium* (Panicoideae: Andropogoneae) to sugarcane. BMC Evolutionary Biology 19:33.

Leseberg CH, Duvall MR. 2009. The complete chloroplast genome of Coix lacryma-jobi and a comparative molecular evolutionary analysis of plastomes in cereals. Journal of Molecular Evolution 69:311-318.

Li S, Qian X, Zheng Z, Shi M, Chang X, Li X, Liu J, Tu T, Zhang D. 2018. DNA barcoding the flowering plants from the tropical coral islands of Xisha (China). *Ecology and evolution*, *8*:10587-10593.

Lloyd Evans D, Joshi SV. 2016. Complete chloroplast genomes of Saccharum spontaneum, Saccharum officinarum and Miscanthus floridulus (Panicoideae:

Andropogoneae) reveal the plastid view on sugarcane origins. Systematics and Biodiversity, 14:548–571.

Ng'uni D, Geleta M, Fatih M, Bryngelsson T. 2010. Phylogenetic analysis of the genus Sorghum based on combined sequence data from cpDNA regions and ITS generate well-supported trees with two major lineages. *Annals of botany*, *105*:471-480.

Orton, L.M., 2015. *Phylogenomic study of selected species within the genus Zea: Mutation rate analysis of complete chloroplast genomes* (Doctoral dissertation, Northern Illinois University).

Saski C, Lee SB, Fjellheim S, Guda C, Jansen RK, Luo H, Tomkins J, Rognli OA, Daniell H, Clarke JL. 2007. Complete chloroplast genome sequences of Hordeum vulgare, Sorghum bicolor and Agrostis stolonifera, and comparative analyses with other grass genomes. Theoretical and Applied Genetics 115:571–590.

Schnable PS, Ware D, Fulton RS, Stein JC, Wei F, Pasternak S, Liang C, Zhang J, Fulton L, Graves TA, Minx P. 2009. The B73 maize genome: complexity, diversity, and dynamics. Science 326:1112–1115.

Skendzic EM, Columbus JT, Cerros-Tlatilpa R. 2007. Phylogenetics of Andropogoneae (Poaceae: Panicoideae) based on nuclear ribosomal internal transcribed spacer and chloroplast trnL–F sequences. *Aliso: A Journal of Systematic and Evolutionary Botany*, *23*:530-544.

Teerawatananon A, Jacobs SW, Hodkinson TR, 2011. Phylogenetics of Panicoideae (Poaceae) based on chloroplast and nuclear DNA sequences. *Telopea*, *13*:115-42.

Thornhill AH, Baldwin BG, Freyman WA, Nosratinia S, Kling MM, Morueta-Holme N, Madsen TP, Ackerly DD, Mishler BD. 2017. Spatial phylogenetics of the native California flora. *BMC biology*, *15*:96.

Tsuruta SI, Ebina M, Kobayashi M, Takahashi W. 2017. Complete Chloroplast Genomes of Erianthus arundinaceus and Miscanthus sinensis: Comparative Genomics and Evolution of the Saccharum Complex. PloS One, 12:e0169992.

Welker CA, Souza-Chies TT, Longhi-Wagner HM, Peichoto MC, McKain MR, Kellogg, EA. 2016. Multilocus phylogeny and phylogenomics of Eriochrysis P. Beauv.(Poaceae–Andropogoneae): Taxonomic implications and evidence of interspecific hybridization. Molecular Phylogenetics and Evolution 99:155–167.
